# Data driven mathematical model of colon cancer progression

**DOI:** 10.1101/2020.11.02.365668

**Authors:** Arkadz Kirshtein, Shaya Akbarinejad, Wenrui Hao, Trang Le, Rachel A. Aronow, Leili Shahriyari

## Abstract

Every colon cancer has its own unique characteristics, and therefore may respond differently to identical treatments. Here, we develop a data driven mathematical model for the interaction network of key components of immune microenvironment in colon cancer. We estimate the relative abundance of each immune cell from gene expression profiles of tumors, and group patients based on their immune patterns. Then we compare the tumor sensitivity and progression in each of these groups of patients, and observe differences in the patterns of tumor growth between the groups. For instance, in tumors with a smaller density of naive macrophages than activated macrophages, a higher activation rate of macrophages leads to an increase in cancer cell density, demonstrating a negative effect of macrophages. Other tumors however, exhibit an opposite trend, showing a positive effect of macrophages in controlling tumor size. Although the results indicate that for all patients, the size of the tumor is sensitive to the parameters related to macrophages such as their activation and death rate, this research demonstrates that no single biomarker could predict the dynamics of tumors.

## 1 Introduction

Recent studies show that many cancers arise from sites of chronic inflammation [1–4]. Balkwill et. al. [5] provide a list of inflammatory conditions that predispose an individual to cancer, in particular to colorectal cancer. Indeed, inflammatory bowel diseases like ulcerative colitis and colonic Crohn’s disease are strongly associated with colorectal cancer [6]. In one experiment, chronic ulcerative colitis was induced in mice and, fourteen weeks later, the mice developed colitis-associated cancer (CAC) [7].

Most common cancer treatments are designed to kill tumor cells. However, the way in which cells die is very important, because dying cells may release molecules that initiate an immune response. We shall refer to cells that go through the process of necrotic cell death as necrotic cells. Necrotic cells are known to release damage-associated molecular pattern (DAMP) molecules such as high mobility group box 1 (HMGB1), which triggers immune responses [8, 9]. In particular, HMGB1 activates dendritic cells [10]. There is an evidence that the expressions of HMGB1 and RAGE, its receptor, are significantly higher in ulcerative colitis than in control cases [11]. HMGB1 has been observed in other cancers, as a result of treatments by radiotherapy and chemotherapy [10, 12–14].

In colon cancer, activated CD8^+^ T cells enhance production of necrotic cells by expressing high levels of cytokines like IFN-*γ* and FasL [15]. Necrotic cells and macrophages release HMGB1 to activate dendritic cells [10], which leads to activation of T-cells [16]. In addition, intestinal epithelial cells, which are in close contact with DCs, activate dendritic cells by releasing molecules like thymic stromal lymphopoietin (TSLP) [17, 18]. Once activated, dendritic cells release cytokines STAT4, STAT6, and IL-4,which induce differentiation of naive T-cells into effector T cells (Th1, Th17, and Th2) [19]. CD4^+^ T-cells can also become activated by TNF-*α*, which is released by *M*_1_ macrophages [20]. Activated CD4^+^ T-cells release IL-2, 4, 5, 13 and 17 to activate killer cells like CD8^+^ T-cells [16, 21, 22]. CD4^+^ T-cells also release IFN-*γ*, which activates *M*_1_ macrophages [23, 24].Activated macrophages and CD4^+^ effector T-cells release tumor-promoting cytokines interleukin 6 (IL-6) [25]. IL-6 promotes tumor growth by activating STAT3 in intestinal epithelial cells [26].

All these observations indicate the importance of the relative abundance of various immune cells, as well as their interaction networks, in the colonic tumors’ initiation and progression. In the present paper, we develop a data driven mathematical model of colon cancer with emphasis on the role of immune cells. We use cancer patients’ data to estimate the percentage of each immune cell types in their primary tumors. The developed mathematical model is based on the network shown in Figure 1, and it is represented by a system of ordinary differential equations (ODEs) within the tumor.

**Figure 1:**
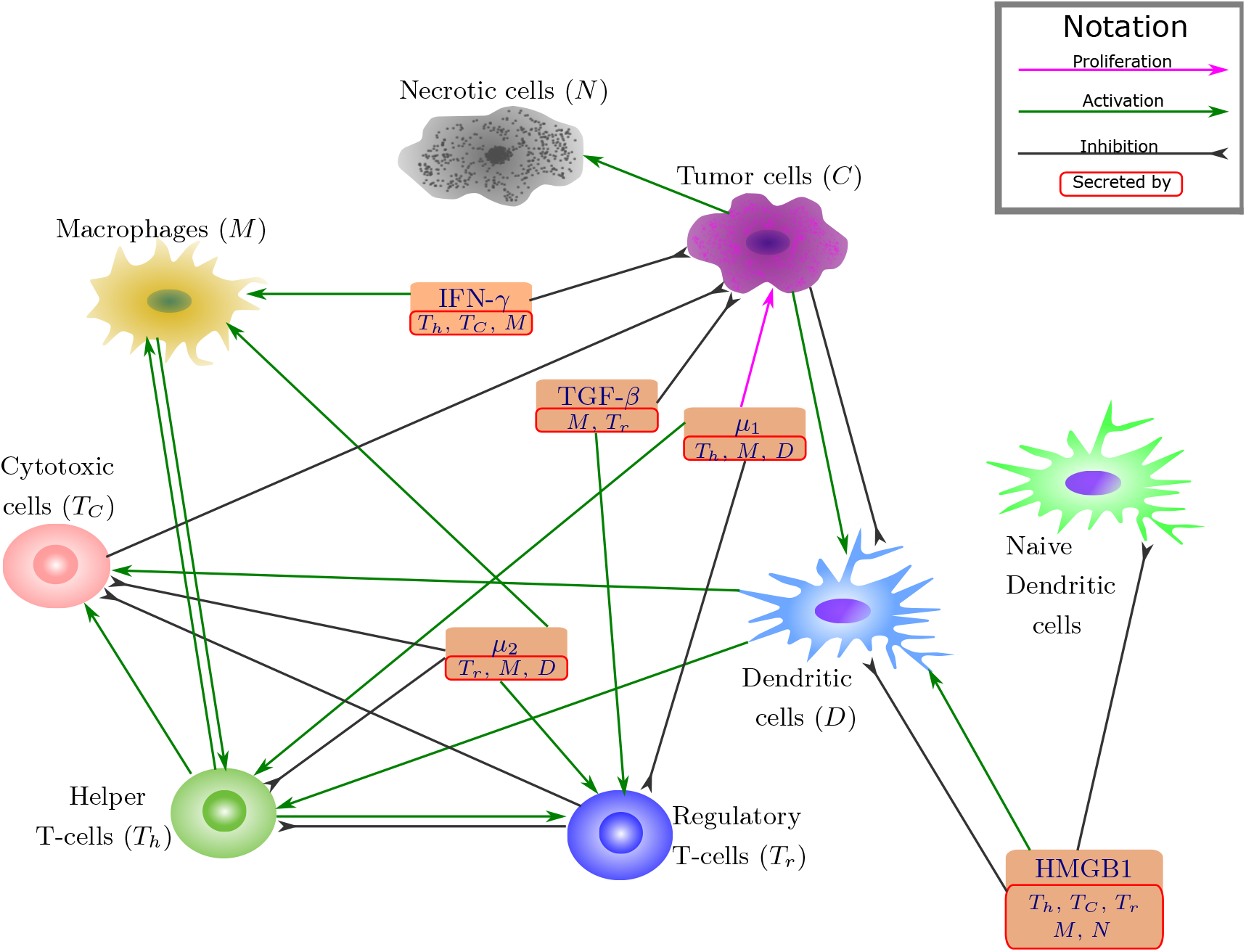
Network of cells and cytokines. Sharp arrows indicate activation or proliferation, and the blucked arrow indicates inhibitions.

## 2 Materials and Methods

### 2.1 Mathematical model

We develop a mathematical model for colon cancer based on the interaction network among key players in colon cancer shown in Figure 1, and the list of variables is given in Table 1. The model is represented by a system of differential equations for concentrations and changing in time in unit of day. For clarity, we develop a simplified model in terms of ordinary differential equations. For biochemical processes *A* + *B → C*, we use the mass action law 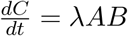, where *λ* is production rate of *C* [27, 28]. Throughout the paper, we use the symbol *λ* for production, activation or proliferation rates, and the symbol *δ* for decay, natural death (apoptosis) or premature death (necrosis) rates.

**Table 1:** Model’s Variables. Names and descriptions of variables used in the model.

| Variable | Name | Description |
| --- | --- | --- |
| $T_N$ | Naive T-cells | |
| $T_h$ | Helper T-cells | |
| $T_C$ | Cytotoxic cells | includes CD8+ T-cells and, possibly, NK cells |
| $T_r$ | Regulatory T-cells | |
| $D_n$ | Naive dendritic cells | |
| $D$ | Activated dendritic cells | antigen presenting cells |
| $M$ | Macrophages | |
| $C$ | Cancer cells | |
| $N$ | Necrotic cells | |
| $H$ | HMGB1 | |
| $\mu_1$ | Carcinogenic cytokines | includes effects of IL-6, IL-17, IL-21 and IL-22 |
| $\mu_2$ | Immunosuppressive agents | includes effects of IL-10 and CCL20 |
| $I_\gamma$ | IFN-γ | |
| $G_\beta$ | TGF-β | |

#### 2.1.1 Cytokine approximation

In order to reduce the complexity of the system, we treat some of cytokines as independent variables and approximate the value of other cytokines through already existing variables. Additionally, we combine the cytokines that have a similar function in the interaction network (Figure 1). So, we combine IL-6, IL-17, IL-21 and IL-22 and denote their sum by the variable *µ*_1_. We also combine IL-10 and CCL20 and denote their sum by the variable *µ*_2_. The cytokines treated as model variables are HMGB1, IFN-*γ*, TGF-*β*, IL-6, and IL-10. We then model the dynamics of cytokines in the following way.

HMGB1 is passively released from necrotic cells [29], or actively secreted from activated T-cells cells and macrophages [30, 31]. Thus we can model the dynamics of HMGB1 by the equation:

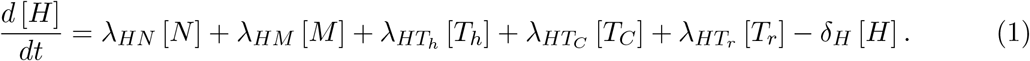

IL-6 is secreted by TAMs [25,32–34], helper T-cells [25,34–36] and sub-population of dendritic cells [37, 38]. IL-17, IL-21 and IL-22 are produced by helper T-cells [39]. So the resulting dynamics for [*µ*_1_] can be written as

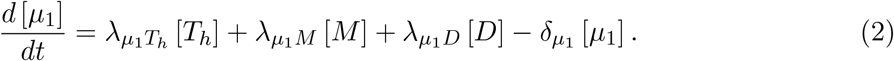

IL-10 is produced by macrophages [40, 41], dendritic cells [37,42] and Treg cells [35,39,43,44]. CCL20 is produced by macrophages [45]. Thus, the equation for [*µ*_2_] is

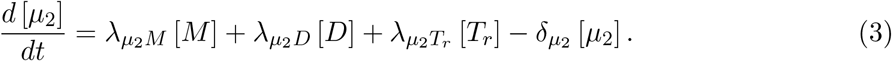

IFN-*γ* is secreted by a sub-population of macrophages [40, 46–49], helper T-cells [23, 24] and cytotoxic cells [15], which results in the following equation:

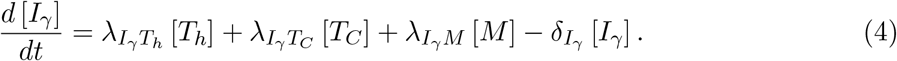

TGF-*β* is produced by macrophages [40, 41] and T-reg cells [35, 39, 43, 50] leading to the equaiton:

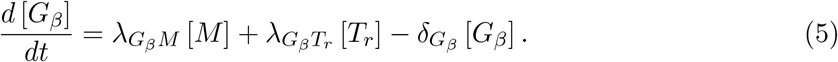

Other cytokines, like IL-2, IL-4, IL-5, and IL-13, we consider to be in a quasi-equilibrium state, i.e. proportional to the concentration of cells that secrete/produce them. In particular, IL-2, IL-5, and IL-13 are produced by CD4+ T-cells [16, 22, 51], so we consider

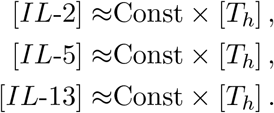

IL-4 is also produced both by CD4+ T-cells [16, 22, 51] and dendritic cells [19], so we take

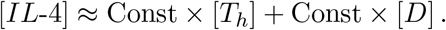

IL-12 secreted by macrophages [40, 41] and dendritic cells [32, 33, 37, 39, 52, 53], thus can be approximated as

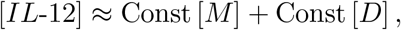

while IL-23 and TNF-*α* are secreted solely by macrophages [32, 33], hence their approximation is

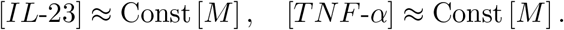

#### 2.1.2 T-cells

In this model we differentiate four subgroups of T-cells: naive, helper, cytotoxic, and regulatory. Naive T-cells, *T*_*N*_, are not necessarily part of tumor micro-environment, as they usually are activated within lymph nodes. However, making activation rates for other types of T-cells proportional to the density of naive cells creates a better controlled system and avoids unlimited exponential growth. Thus, we summarize the equation for the dynamics of the naive T-cells after detailing the equations of other types of T-cells.

Helper T-cells can be activated with antigen presentation by dendritic cells [16]. CD4+ T-cells can be additionally activated by IL-12, while Th17 are activated by IL-6, TNF-*α*, and IL-23 [39]. Regulatory T-cells inhibit protective immune response (helper and cytotoxic T-cells) in several ways including production of immunosuppresive cytokines such as IL-10 and CCL20 as well as through contact-dependent mechanisms [39]. Additionally, we introduce the apoptosis rate for helper cells 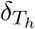 The resulting equation is

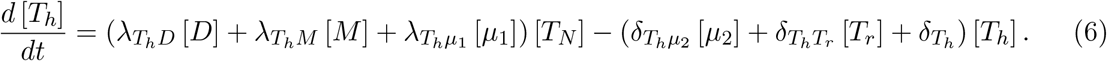

The variable corresponding to cytotoxic cells accounts for the effects of cytotoxic T-lymphocytes (mainly CD8+ T-cells) and possibly natural killer cells. CD8+ T-cells are activated by IL-2, IL-4, IL-5, and IL-13 [16, 22, 51]. Cumulative effect of these cytokines can be written as

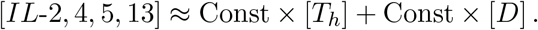

Activation of natural killer cells requires IL-2 [54], which is already included. We also include inhibitory effects mediated by T-reg cells. The dynamics of *T*_*C*_ cell group is modeled by the following equation:

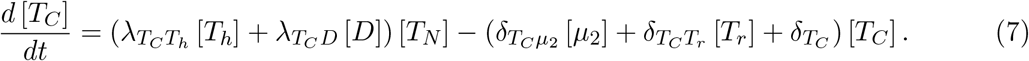

Regulatory T-cells can be activated by IL-2 [39, 55, 56], CCL20 [45] and TGF-*β* [39, 50]. IL-6 suppresses T-reg differentiation and shifts it towards T-helper type [57]. The resulting dynamics can be described as follow:

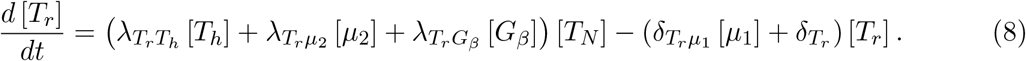

Combining all activation and introducing independent naive T-cell production 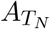, we get the following equation for naive T-cells:

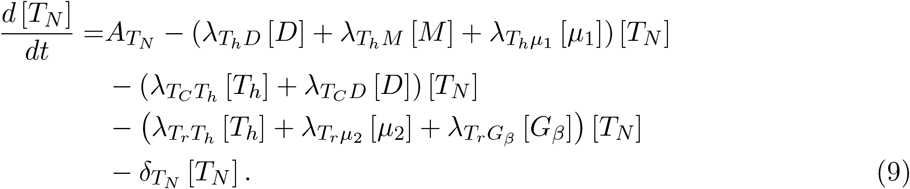

#### 2.3.1 Dendritic cells

Dendritic cells become activated by HMGB1 [10] and TSLP, which is released by epithelial cells [17, 18]. We take TSLP in quasi-equilibrium state as

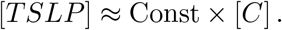

On the other hand multiple factors induced by cancer cells may promote apoptosis of dendritic cells [58–62]. Additionally, there’s evidence that HMGB1 can reduce the maturation rate of dendritic cells [44, 62]. Introducing the independent production rate of naive dendritic cells 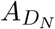 we get the following system for dynamics of naive (*D*_*N*_) and activated (*D*) dendritic cells:

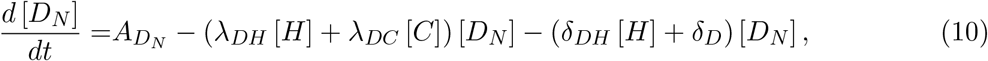

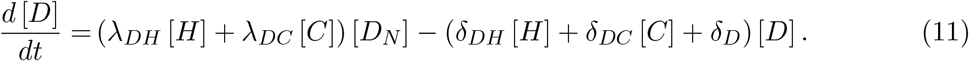

#### 2.1.4 Macrophages

There are two main sub-types of macrophages: M1 and M2. M1 phenotype can be activated by IFN-*γ*, while M2 can be activated IL-4 and IL-13, which are secreted by helper T-cells [40, 41]. Additionally there’s a possibility of tumor associated macrophage (TAM) activation by IL-10 [40, 63, 64]. Introducing naive (*M*_*N*_) and activated (*M*) TAMs, as well as production rate for naive macrophages *A*_*M*_, we can write the following system:

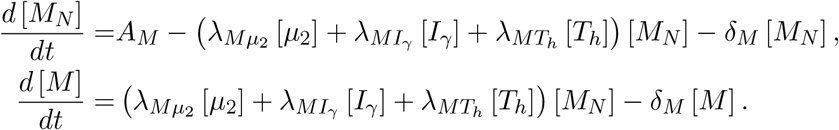

Next, to simplify the system we introduce the total amount of macrophages *M*_0_ = [*M*_*N*_]+[*M*]. Adding the above equations we get 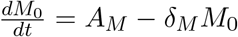 If we assume initial conditions for *M*_0_ to be at the equilibrium *M*_0_ = *A*_*M*_ */δ*_*M*_, then *M*_0_ will remain constant at all times. Then we can express naive macrophages as [*M*_*N*_] = *M*_0_ *−* [*M*] and write the resulting equation for macrophages as follows:

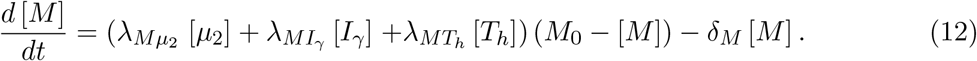

#### 2.1.5 Cancer cells

Cancer cells are epithelial cells with abnormally high growth and abnormally small death rate (apoptosis). Additional loss of apoptosis in cancer cells is induced by IL-6 [58, 60, 65, 66]. In addition to innate abnormally high proliferation rate *λ*_*C*_, proliferation in cancer can be stimulated by expression of STAT3 in cancer cells, where STAT3 is activated by cytokines such as IL-6, IL-17, IL-21, and IL-22 [39, 67]. On the other hand, cancer development is suppressed by TGF-*β* [39, 68–70], IL-12 and IFN-*γ* [39]; the suppressive properties of IL-12 are mediated by IFN-*γ* [71] (so it is not directly included in the equation). Cytotoxic T-cells also directly target cancer cells for destruction [39]. In cancer modeling, proliferation is traditionally taken to be proportional to [*C*] (1 *−* [*C*] */C*_0_), where *C*_0_ is the total capacity [72, 73]. Thus the resulting equation is

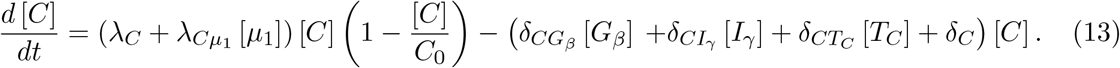

#### 2.1.6 Necrotic cells

We designate cells which go through the process of necrotic cell death as necrotic cells. Since there is a limited amount of resources in the tumor microenvironment, and cells are under pressure, there are always some necrotic cells produced by the tumor. In addition, when activated cytotoxic T-cells kill colorectal cancer cells by expressing high levels of cytokines like IFN-*γ* and FasL [15], a fraction of the cancer cells may go through the stage of first becoming necrotic cells. Therefore, the rate of “production” of the necrotic cells is given by the fraction of dying cancer cells, and the resulting dynamics can be written as follows:

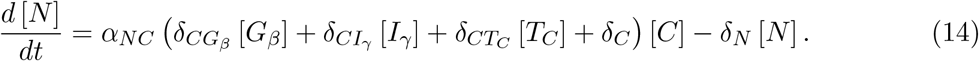

### 2.2 Non-dimensionalization and sensitivity analysis

For additional numerical stability and to eliminate scale dependence, we perform non-dimensionalization of the system. For variable *X* converging to a steady state *X*_*∞*_, we consider non-dimensional variable 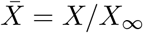 Then, 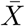 satisfies the equation

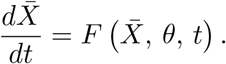

The (first order) solution sensitivity *S* with respect to the model parameter 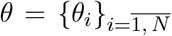 is defined as a vector

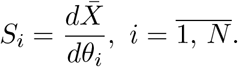

In general, the sensitivity vector is time dependent and varies for different solutions and parameter sets [74–76]. However, here we consider sensitivity at the steady state of the equation. The sensitivity of each parameter in the neighborhood of a chosen parameter set Ω(*θ*) is defined as

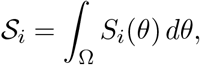

where the integration is evaluated numerically with sparse grid points [77, 78].

We choose three quantities of interest for the sensitivity analysis: amount of cancer cells *C*, total amount of cells, and a measure of how fast the system is converging to the steady state. Consider general steady state system as follows

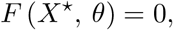

where *X*^*\**^ is the equilibrium. We then consider a small perturbation to *X*^*\**^ as 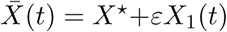. The linearized system becomes

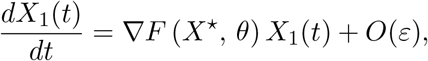

where *∇F* (*X, θ*) is the Jacobian matrix of *F* (*X, θ*) with respect to *X*. Thus we have *X*_1_(*t*) ≈ *e*^*∇F* (*X*\*, *θ*)*t*^ and the minimal eigenvalue min *λ* (*∇F* (*X*^*\**^, *θ*)) determines how fast it reaches the steady state.

### 2.3 Cancer patients’ data

In recent years, several tumor deconvolution methods have been developed to estimate the relative abundance of various cell types in a tumor from its gene expression profile. A review of these methods [79] and an application of CIBERSORTx on renal cancer [80] show a great performance of CIBERSORTx model. To identify the immune profiles of colonic tumors, we applied CIBERSORTx [81] on RNA-seq gene expression profiles of primary tumors of patients with colon cancer from the TCGA project of COAD downloaded from UCSC Xena web portal. There are a total of 329 patients with RSEM normalized RNA-seq data in log_2_ scale. Before applying CIBERSORTx on this dataset, we transformed the gene expression values to the linear space.

### 2.4 Numerical methods

In order to solve the time dependent system, we employ the SciPy odeint function [82] using initial conditions based on patients with the smallest tumor area within each cluster. The sensitivity analysis of the system based on the cancer and total cell density at steady state is obtained analytically by differentiating the steady state equation with respect to the parameters, namely,

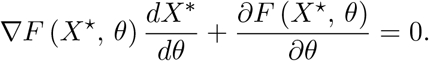

Then to obtain the sensitivity, 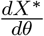, one just needs to numerically invert the matrix *∇F*. On the other hand, it is hard to analytically obtain the sensitivity of the eigenvalue, so instead a finite-difference approach is used as follows:

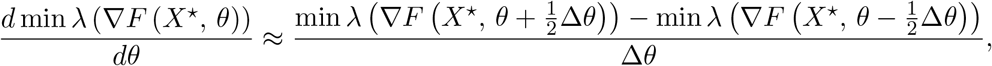

where Δ*θ* is a small discretization parameter.

## 3 Results

We derived an ODE system describing complex dynamics in the colon cancer microenvironment. Assuming non-negative values of all parameters and non-negative initial conditions, the solution of the system remains non-negative and globally bounded (see Appendix A).

### 3.1 Patient data analysis

We downloaded TCGA clinical data, which includes tumor dimension, stage, age at diagnosis, and gene expression profiles of primary tumors for patients with colon cancer from GDC portal. We applied CIBERSORTx B-mode on gene expression profiles to estimate the fraction of each immune cell type in each tumor. Elbow method applied on estimated cell fractions (Figure 2-A) showed the existence of five distinct immune patterns. We hence performed K-means clustering with *K* = 5, to group patients based on the immune pattern of their primary tumors. Figure 2-B shows average cell fractions for patients divided into five clusters based on their immune profiles. To investigate the effect of these immune patterns on the dynamics of tumors, we model each cluster separately, and based on the steady state assumptions (see Appendix B), we generate a parameter set for each cluster, with steady state values derived from patient data as described further.

**Figure 2:**
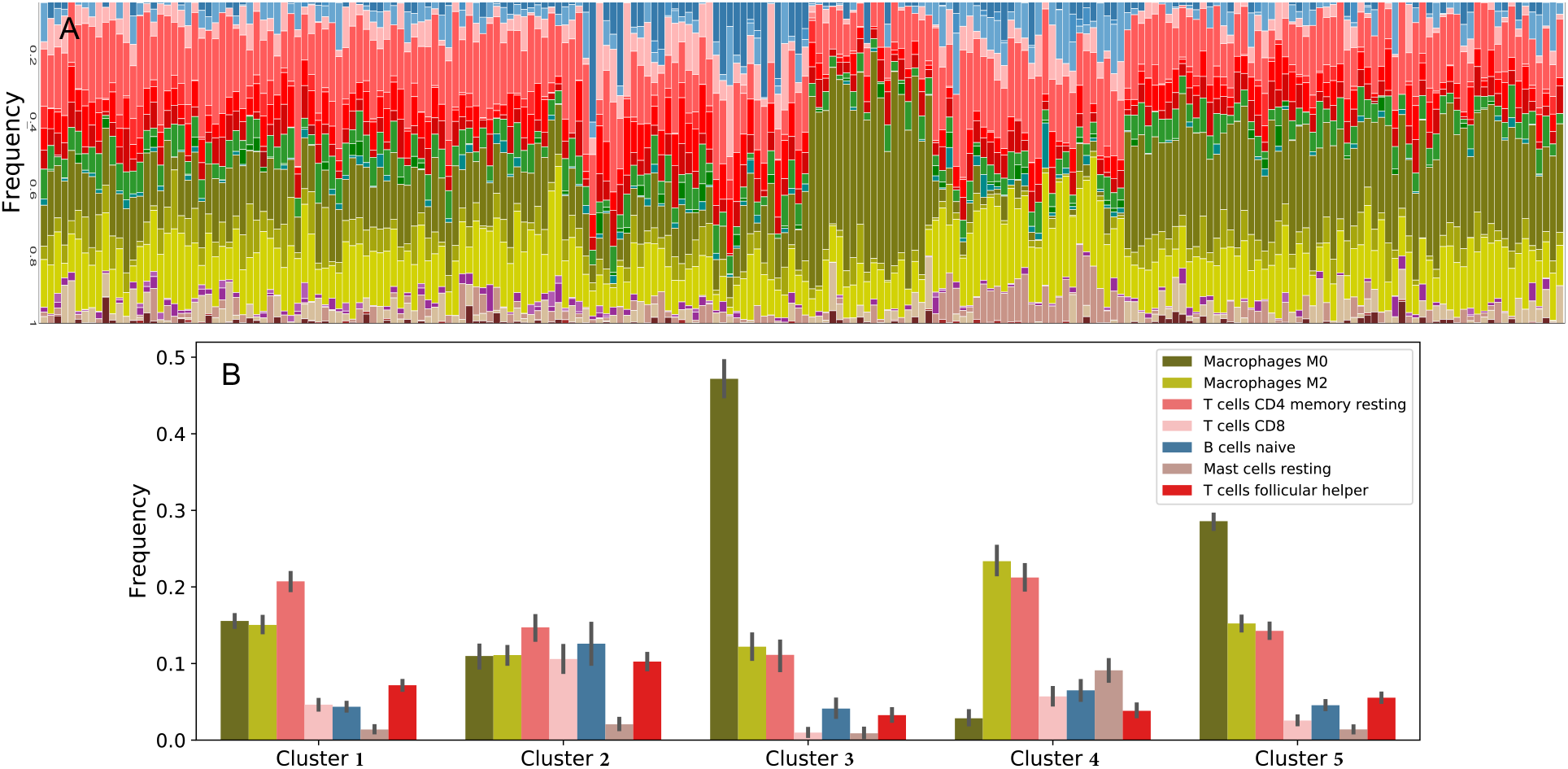
Immune cell fractions. Sub-figure A shows the fraction of immune cells in each colonic tumor. Sub-figure B indicates the frequencies of immune cell types in each cluster of patients. Clusters were formed based on variations of 22 immune cell types, some of which were later combined and others were not included in the model. Cell frequencies on this figure are average within the cluster and vertical bars show the standard deviations

The deconvolution data, described in section 2.3, only provides the ratios of immune cells in the tumor microenvironment. For each patient *P*, we define their size of tumor (size(*P*)) to be the product of the longest and the shortest dimensions of the tumor, and we assume total cell density is proportional to the size of the tumor:

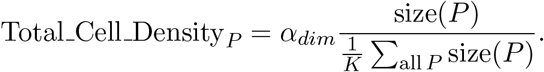

Then, we take each immune cell value from deconvolution multiplied by 0.4*α*_*dim*_ Σ(Immune cell ratios) and

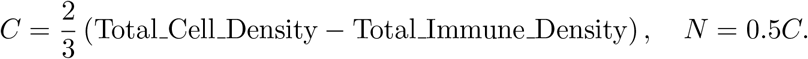

For each cluster, we consider the mean of variables of patients with tumor size above the average of their cluster as the steady state values of the variables for the corresponding cluster. The resulting data is given in Table 2.

**Table 2:** Steady state cell densities. Mean cell densities in cells/cm^3^ for each cluster are used to derive parameter sets for sensitivity analysis and dynamics computations. Calculated based only on patients with tumor size above average for each cluster.

| Cluster | $T_N^\infty$ | $T_h^\infty$ | $T_C^\infty$ | $T_r^\infty$ | $D_N^\infty$ | $D^\infty$ | $M^\infty$ | $M_0$ |
| --- | --- | --- | --- | --- | --- | --- | --- | --- |
| 1 | 1.4914e+4 | 4.6358e+3 | 2.5845e+3 | 2.3891e+3 | 3.0504e+2 | 6.0214e+2 | 1.1798e+4 | 2.1004e+4 |
| 2 | 1.1429e+4 | 6.0411e+3 | 5.3853e+3 | 3.3646e+3 | 1.0329e+2 | 5.1299e+2 | 8.6227e+3 | 1.6445e+4 |
| 3 | 9.2381e+3 | 1.3864e+3 | 1.1139e+3 | 2.7910e+3 | 1.8878e-1 | 1.8635e+2 | 6.7972e+3 | 3.2146e+4 |
| 4 | 1.3878e+4 | 2.4910e+3 | 3.2172e+3 | 2.2783e+3 | 1.4196e+2 | 6.2154e+2 | 1.2931e+4 | 1.5761e+4 |
| 5 | 1.0262e+4 | 3.7844e+3 | 1.6853e+3 | 2.6394e+3 | 8.0199e+1 | 1.9084e+2 | 1.1603e+4 | 2.8198e+4 |
| | $C^\infty$ | $N^\infty$ | $\mu_1^\infty$ | $\mu_2^\infty$ | $H^\infty$ | $I_\gamma^\infty$ | $G_\beta^\infty$ | |
| 1 | 9.1531e+4 | 4.5765e+4 | 1.6328e+2 | 1.2987e+3 | 8.9811e+3 | 8.5737 | 1.9037e+4 |  |
| 2 | 9.7064e+4 | 4.8532e+4 | 1.7552e+2 | 1.3249e+3 | 8.5279e+3 | 10.5677 | 2.2275e+4 |  |
| 3 | 9.0029e+4 | 4.5014e+4 | 1.9866e+2 | 1.2906e+3 | 9.5122e+3 | 0.8287 | 2.5145e+4 |  |
| 4 | 9.6956e+4 | 4.8478e+4 | 1.1410e+2 | 3.3689e+2 | 5.1782e+3 | 1.2703 | 8.1734e+3 |  |
| 5 | 8.0584e+4 | 4.0292e+4 | 1.2058e+2 | 7.1551e+2 | 7.7848e+3 | 5.7892 | 2.6260e+4 |  |

While macrophage capacity *M*_0_ is derived from the data, we assume cancer capacity to be *C*_0_ = 2*∗ C* for both mean-based and extreme-based data. We choose *α*_*dim*_ = 1.125e+05 to approximately match the average density of cancer cells across all patients to 4.5e+04 cells/cm^3^ reported in [83]. However, it is important to note that this is no more than scaling and has no effect on the dynamics of the dimensionless system.

For the time-dependent calculations, we choose initial conditions for each cluster based on patients with the smallest tumor size. The relative values are given in Table 3. The dynamics with initial conditions based on other patients is presented in Appendix C.

**Table 3:**
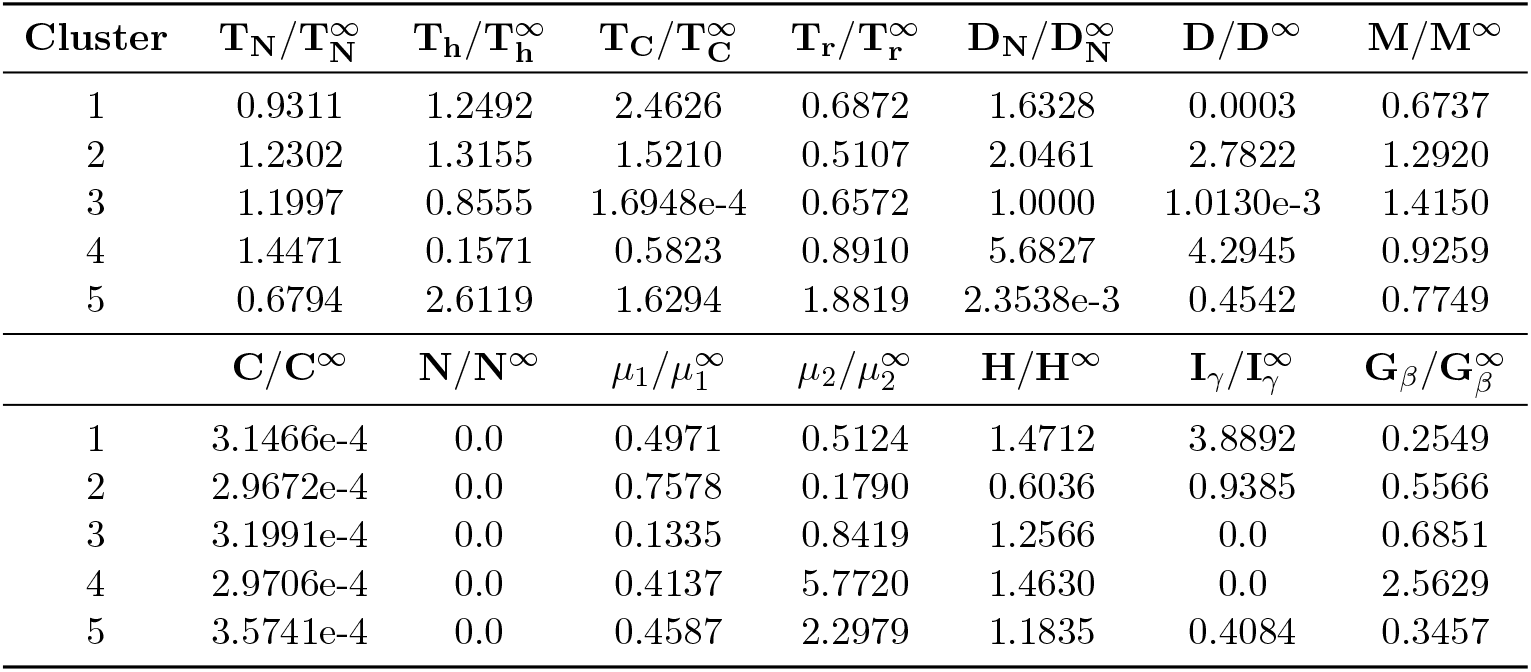
Dimensionless initial conditions. Values of initial conditions for the dimensionless system derived from the patients with the smallest tumor size.

| Cluster | $T_N/T_N^\infty$ | $T_h/T_h^\infty$ | $T_C/T_C^\infty$ | $T_r/T_r^\infty$ | $D_N/D_N^\infty$ | $D/D^\infty$ | $M/M^\infty$ |
| --- | --- | --- | --- | --- | --- | --- | --- |
| 1 | 0.9311 | 1.2492 | 2.4626 | 0.6872 | 1.6328 | 0.0003 | 0.6737 |
| 2 | 1.2302 | 1.3155 | 1.5210 | 0.5107 | 2.0461 | 2.7822 | 1.2920 |
| 3 | 1.1997 | 0.8555 | 1.6948e-4 | 0.6572 | 1.0000 | 1.0130e-3 | 1.4150 |
| 4 | 1.4471 | 0.1571 | 0.5823 | 0.8910 | 5.6827 | 4.2945 | 0.9259 |
| 5 | 0.6794 | 2.6119 | 1.6294 | 1.8819 | 2.3538e-3 | 0.4542 | 0.7749 |
| | $C/C^\infty$ | $N/N^\infty$ | $\mu_1/\mu_1^\infty$ | $\mu_2/\mu_2^\infty$ | $H/H^\infty$ | $I_\gamma/I_\gamma^\infty$ | $G_\beta/G_\beta^\infty$ |
| 1 | 3.1466e-4 | 0.0 | 0.4971 | 0.5124 | 1.4712 | 3.8892 | 0.2549 |
| 2 | 2.9672e-4 | 0.0 | 0.7578 | 0.1790 | 0.6036 | 0.9385 | 0.5566 |
| 3 | 3.1991e-4 | 0.0 | 0.1335 | 0.8419 | 1.2566 | 0.0 | 0.6851 |
| 4 | 2.9706e-4 | 0.0 | 0.4137 | 5.7720 | 1.4630 | 0.0 | 2.5629 |
| 5 | 3.5741e-4 | 0.0 | 0.4587 | 2.2979 | 1.1835 | 0.4084 | 0.3457 |

### 3.2 Sensitivity analysis

We perform sensitivity analysis of the non-dimensionalized system with parameters derived from patient data through steady state assumptions. Table 2 contains the steady state values used for each cluster, and Appendix B shows the parameter derivation and non-dimensionalization in detail. We use cancer cells, total cell density and minimal eigenvalue of the Jacobian of the ODE system as the variables of interest in the sensitivity analysis. Minimal eigenvalue of the Jacobian serves as a measure of how fast the system converges to the steady state. Figure 3-A shows the four most sensitive parameters for each cluster. Additionally, to evaluate the effect of immune microenvironment on cancer, we look at the sensitivity of cancer cells and total cell density excluding the parameters appearing in the equations for cancer and necrotic cells. The resulting data denoted as “Immune sensitivity” is given in Figure 3-B.

**Figure 3:**
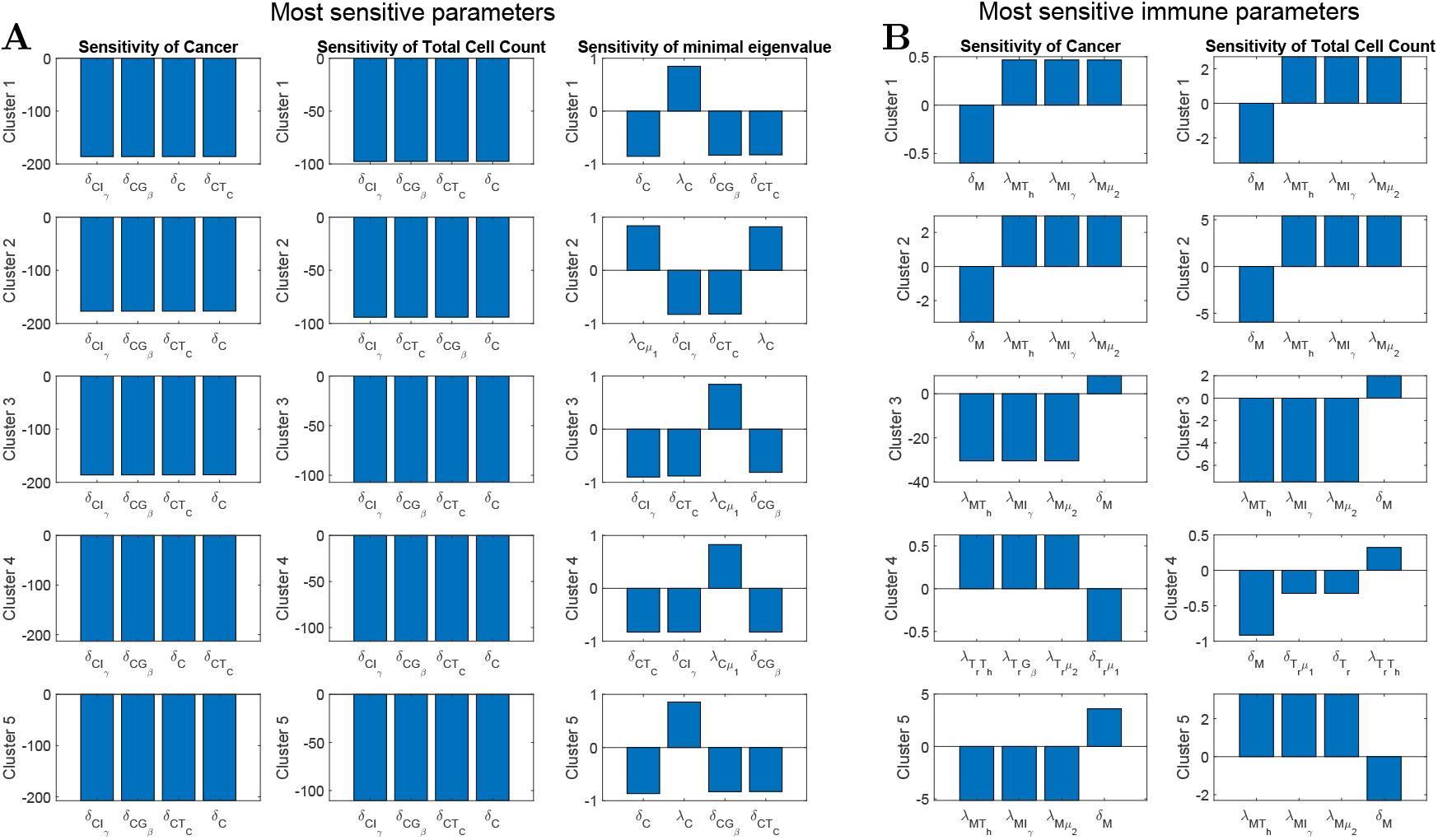
Sensitivity analysis. The first, second, and third columns of sub-figure A respectively present the results of non-dimensional sensitivity of cancer cell density, total cell density and minimal eigenvalue of the Jacobian of the system at the steady state. Minimal eigenvalue is used as a measure of how fast the system converges to the steady state. Sub-figure B shows the sensitive parameters related to immune cells. Each row of plots shows the most sensitive parameters for each cluster of patients.

Across all clusters the most sensitive parameters are cancer proliferation and death rates directly present in the cancer equation (13). From third column in Figure 3-A, we conclude that for all clusters increased cancer proliferation coefficients correspond to faster convergence to the steady state, while increased cancer death rates lead to a slower convergence. When considering immune sensitivity presented on figure 3-B, in clusters 1, 2, 3, and 5, the most sensitive immune parameters are those corresponding to the activation and decay rates of macrophages, with only sensitivity levels being different between clusters and variables. In clusters 1 and 2, which include tumors with a smaller density of naive macrophages than activated macrophages, an increase in decay rate of macrophages causes a decrease in the density of cancer cells and total cell density. On the other hand, an increase in any of the activation rates for macrophages causes an increase in both quantities of interest. However, for clusters 3 and 5, which include tumors with a higher density of naive macrophages than activated macrophages, the effects are reversed. Interestingly, for cluster 3, the increase in macrophage activation rate results in both lower cancer cell density and total cell density, with latter sensitivity being noticeably smaller by absolute value. On the other hand, for cluster 5 the increase in macrophage activation rate results in lower cancer cell density, but higher total cell density. This can be explained by a significant increase in immune cell density, which for cluster 5 is even higher than the corresponding decrease in cancer cell density. All these results demonstrate that at the steady state tumor-associated macrophages could have different effects on different clusters of patients depending on their immune profile.

The outlying cluster 4, which consists of tumors with a significantly small density of naive macrophages compared to the other clusters, is less sensitive to the activation rates of macrophages. The most sensitive immune parameters for cancer cell density are those related to the activation and degradation of regulatory T-cells. The results indicate that increased regulatory response activation rate corresponds to an increase in the cancer cell density, while an increase in T-reg cell degradation rate results in a decrease in cancer cell density, demonstrating that for this cluster of patients regulatory T-cells have mostly negative effect. Importantly, the most sensitive parameter for the total cell density is still the decay rate of macrophages, and macrophages still have a negative effect; i.e. the faster decay of macrophases leads to the smaller tumors in the steady state.

### 3.3 Dynamic of tumor microenvironment

We investigate the dynamics of each variable, with parameters derived for each cluster based on steady state assumptions (see Table 2 for steady state values and Tables A1-A3 for parameter values) and initial conditions of patients with the smallest tumor (see Table 3). Figures 4 and 5 are respectively show the dynamics of cell densities and cytokines expressions.

**Figure 4:**
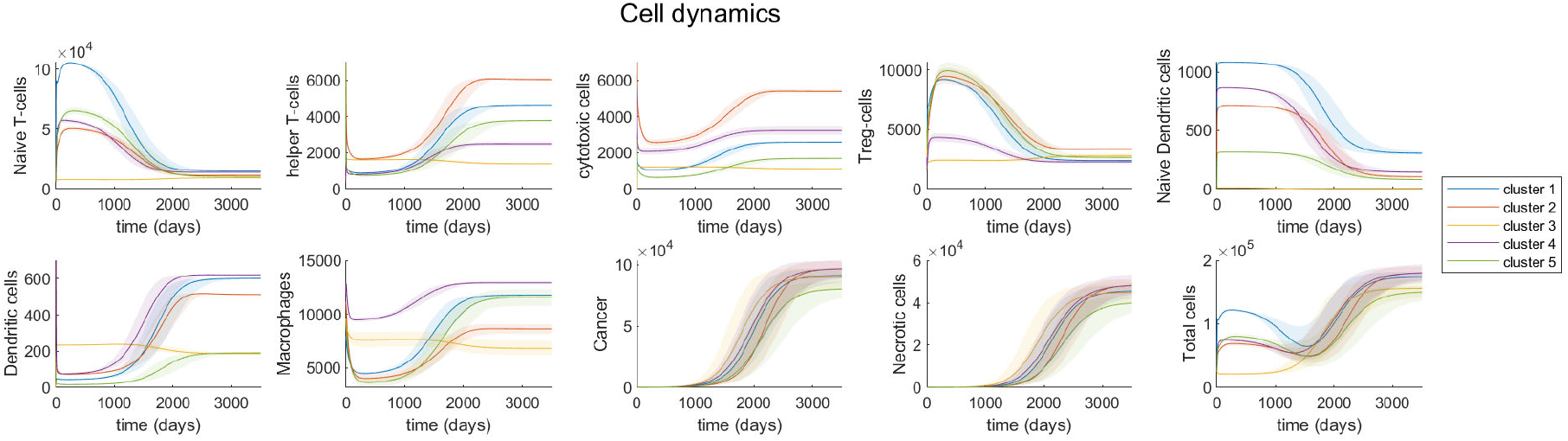
Cells’ dynamics in colonic tumors. Time evolution of cells’ density (cell/cm^3^) for each cell type in the model and total cell density. Different colors represent the models derived for different clusters of patients and shaded regions represent the 10% variation in the most sensitive parameters.

**Figure 5:**
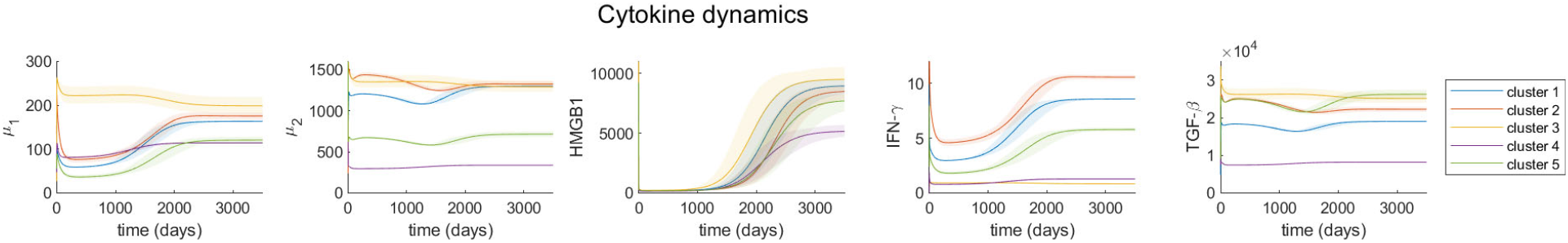
Cytokines’ dynamics in colonic tumors. Time evolution of RNA-seq expression rate of cytokines. Different colors represent the models derived from different clusters of patients and shaded regions represent the 10% variation in the most sensitive parameters.

For most clusters, cancer cells grow as helper T-cells, cytotoxic cells (cytotoxic T-cells and NK cells), dendritic cells and macrophages increase in density over time, while naive T-cells, regulatory T-cells and naive dendritic cells decrease in density. The increase in cytotoxic cells along with tumor progression is somewhat contradicting to the finding in [84, 85] that colon primary tumor growth is associated with decreased cytotoxic T-cells density. However, there is no correlation between tumor size and cytotoxic cells in the TCGA data of colonic primary tumors. Moreover, it is important to note that in our model cancer cells’ growth is multiple times faster than the rate of change of any immune cells (Figure 4). Thus, even though cytotoxic cells density grows over time, the tumor is growing at a much faster rate. Since tumor cells activate dendritic cells which then activate cytotoxic cells, it is reasonable to see some growth of cytotoxic cells when tumor cells density increases rapidly.

Cluster 2 and 4 have the highest cancer cell density at steady state and also the highest growth rate of cancer cells. Cluster 2’s cancer cells start out with lowest growth rate, but at around 1,800 days grow significantly faster and end up growing the fastest among all clusters. Cluster 2 has the highest density of helper T-cells and cytotoxic cells, both in the early stages of cancer development and at steady state, as well as the highest growth rate of these cells. However, cluster 2 has rather low density and low growth rate of macrophages.

Cluster 4, having the largest density of activated macrophages and a significantly small density of naive macrophages (Figure 2-B), first demonstrates average cancer growth rate, but then increases and has one of the 2 highest cancer cell densities at the steady state (Figure 4). Similar to cluster 2, cluster 4 has high density of cytotoxic cells (CD8 T-cells and NK cells) initially and at steady state. Both cluster 2 and 4 have low growth rate of macrophages and high density of dendritic cells, compared to other clusters. Immune cell dynamics of cluster 2 and 4 demonstrate that high density of cytotoxic cells and dendritic cells, along with low growth rate of macrophages correlate with high growth rate of cancer cells.

However, unlike cluster 2, cluster 4 has low growth rate of cytotoxic cells and helper T-cells, and low density of helper T-cells overall. Though both cluster 2 and 4 have low growth rate of macrophages, cluster 4 has the highest density of macrophages among all clusters, while cluster 2 has the second lowest macrophages density. Regulatory T-cells also behave very differently between cluster 2 and cluster 4. Cluster 2 has high density and high decline rate of regulatory T-cells over time, but cluster 4 has both low density and low decline rate of this cell. These observations suggest that cell densities alone cannot predict cancer progression and there are no specific biomarkers that are sufficient to model tumor growth. Instead, a time series immune interaction network with tumor cells can be useful in modeling cancer development.

Cluster 5, with the density of activated macrophages being slightly less than naive macrophages (Figure 2-B), has the lowest cancer cell density at steady state and the lowest cancer cell growth of all clusters (Figure 4). This cluster has the lowest growth rate and density at initial condition and steady state of naive dendritic cells, activated dendritic cells and cytotoxic cells, except for cytotoxic cells density at steady state (second lowest). It also has the highest growth rate of macrophages among the five clusters. This observation might imply that slow tumor growth is associated with low density and growth rate of naive and activated dendritic cells, cytotoxic cells and high growth rate of macrophages.

Cluster 1, which is characterized by the second largest population of macrophages and helper T-cells, demonstrates that dendritic cells alone cannot be chosen as a marker of cancer progression, as it has the second highest dendritic cell population, but only third highest cancer population at the steady state, being surpassed by cluster 2.

Cluster 3, being a clear outlier in the immune dynamics, has near zero density of naive dendritic cells. This alone prevents it from creating significant variations in the immune response during the cancer progression. It is interesting to note, that while almost unchecked by immune responses, this cluster initially demonstrates noticeably highest cancer growth rate, but results in the second lowest cancer density at the steady state.

Tumor cytokines’ dynamics (Figure 5) indicate that as tumor grows, HMGB1, IFN-*γ* and *µ*_1_ (IL-6, IL-17, IL-21, IL-22) increase in density, but TGF-*β* and *µ*_2_ (IL-10, CCL20) stay relatively constant. Cluster 2 and 4, which have the highest cancer cell growth rate among all clusters, show different cytokines’ behaviors throughout time. At steady state, cluster 4 has significantly lower densities of all cytokines in our model than cluster 2, despite the fact that they have the same cancer cell density then. Cluster 4 also has much lower growth rate of *µ*_1_, HMGB1 and IFN-*γ* compared to cluster 2. Cluster 1 and 5 have more similar growth rate of these cytokines as cluster 2, even though they have rather different tumor growth rate from cluster 2. Thus, the density or growth of any specific cytokine is not an adequate predictor of tumor progression, and we need the full interaction network to effectively model the cancer cell growth.

Additionally within each cluster we look at the dynamics of cancer and total cell density with different initial conditions, each derived from a different patient in that cluster. See Appendix C, and specifically figures A1-A5, for more details on different initial conditions and resulting dynamics. This result indicates that even within the same cluster different initial immune profile may cause dramatic difference in cancer progression rate. Additionally, while the dynamics of cancer cell density remains monotone across all patients, we observe oscillatory behavior in the total cell density. This can be explained by a temporary surge of immune cell density at the early stages of cancer, which also appears to correlate with slower cancer progression rate. The only cluster which does not exhibit this oscillatory behavior is cluster 3. As mentioned before, due to lack of naive dendritic cells cluster 3 does not show significant immune cell density variations, which are the source of oscillatory behavior for other clusters.

## 4 Discussion

There are many mathematical models for cancer [86–106]. The approach of many of these mathematical models is varying the parameters values and initial conditions to investigate their effects on the dynamics. However, new advances in tumor deconvolution techniques help us to utilize cancer patients’ data in order to develop a data driven mathematical model of tumor growth. Using tumor deconvolution methods, we estimate the relative abundance of various cell types from gene expression profiles of tumors. The machine learning algorithm of K-means clustering indicates the existence of five distinct groups of colon cancers based on their immune patterns. The comparison of tumor behaviours in these groups suggests that the dynamics of tumors strongly depends on their immune structure.

While it would be ideal to use time course gene expression data of colon cancer patients in our framework, the availability of these time series data sets is limited. In order to combat this limitation, clustering was used to group patients with similar immune patterns and treat each group as time course data based on the size of tumor, which means the data points with small tumor density are considered data from early stages and the data points with large tumor density are considered data from late stages. This method of artificially creating time course data is based on the assumption that immune variation between clusters of patients at any time point is greater than the immune variation within one cluster during tumor progression.

The mathematical model with these assumptions indicate that high density of cytotoxic T-cells and dendritic cells and low growth rate of macrophages are associated with high growth rate of cancer cells, while low density and growth rate of naive and active dendritic cells, cytotoxic T-cells and high growth rate of macrophages correlate with slow tumor growth. In particular, our results imply that macrophages’ growth rate is negatively correlated with tumor growth rate, which is consistent with the observation that high level of macrophages is associated with favorable outcome of colon cancer patients in [84]. This study [84] also shows that high level of regulatory T-cells is related to poor prognosis of patients, which supports our results that regulatory T-cells decrease in density as cancer cells increase in density. Another similar finding between [84] and our study is that the density of dendritic cells increases along with tumor progression.

There is a significant body of research analyzing statistical and mathematical relations of particular components of tumor microenvironment and the disease progression and outcome for subsequent establishment of prognostic biomarkers [83, 107–115]. Our result demonstrates that the dynamics of cancer development cannot be captured by one specific biomarker, but can rather be characterized by complex time-dependent interactions between many components of the immune system and tumor tissue. It is important to further develop and analyze these tumor-immune cell interactions and how they affect different possibilities of treatment.

One way forward is the design of patient-specific models [116–119]. These models can utilize the tumor immune microenvironment deconvolution and clustering methods for available patient data as detailed in this paper. New prognosis can be built based on established dynamics from patients with similar immune characteristics. To better match the dynamics of the model to real patient data, various parameter fitting algorithms can be utilized [120–123]. Another possible improvement is a transition to a partial differential equations model [124] to analyze spatial properties of tumor development as well as temporal.

## Author Contributions

conceptualization, L.S.; methodology, A.K., L.S., W.H., S.A.; software, A.K.; validation, A.K.; formal analysis, A.K., T.L., R.A.; investigation, A.K.; resources, S.A., T.L., R.A.; data curation, S.A., T.L., R.A.; writing–original draft preparation, A.K., L.S., S.A., W.H., T.L., R.A.; visualization, A.K., S.A., W.H.; supervision, L.S.; project administration, L.S.; funding acquisition, L.S.

## Data and Code Availability

Publicly available TCGA clinical data was downloaded from GDC portal. Figure 2 was obtained using TumorDecon software: https://github.com/ShahriyariLab/TumorDecon. Python scripts for computations and MatLab scripts for plotting the results presented on figures 3-5 and A1-A5 are available here: https://github.com/ShahriyariLab/Data-driven-mathematical-model-for-colon-cancer.

## Abbreviations

The following abbreviations are used in this manuscript:

CAC: colitis-associated cancer
CCL20: chemokine (C-C motif) ligand 20
COAD: colon adenocarcinoma
DAMP: damage-associated molecular pattern
DCs: dendritic cells
FasL: fas ligand
GEP: gene expression profiles
HMGB1: high mobility group box 1
IFN: interferon
IL: interleukin
NF-*κ*B: nuclear factor kappa B
NK: cells natural killer cells
ODE: ordinary differential equation
RAGE: receptor for advanced glycation endproducts
RNA-seq: ribonucleic acid sequencing
STAT: signal transducer and activator of transcription
TAM: tumor associated macrophage
TCGA: the cancer genome atlas
TGF: transforming growth factor
TNF: tumor necrosis factor
TSLP: thymic stromal lymphopoietin

## Appendix A ODE system and analysis

Combining equations (1)-(14) we obtain the following system

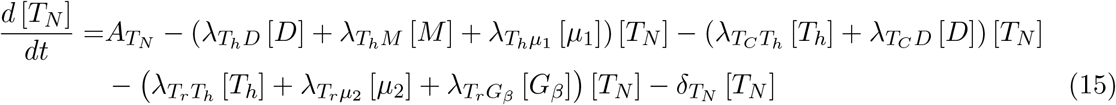

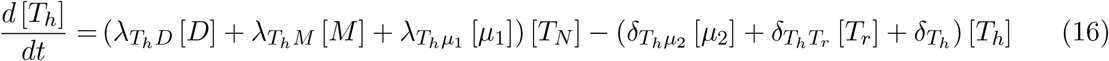

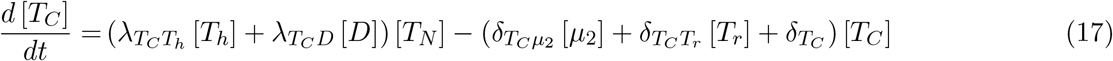

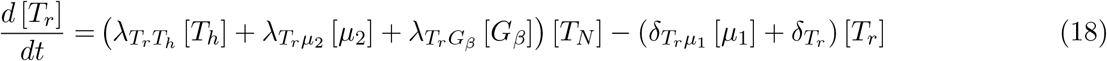

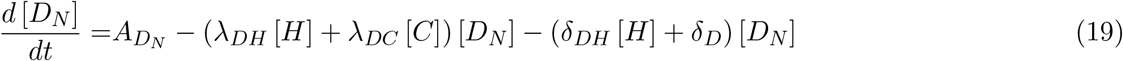

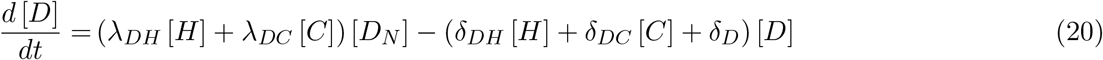

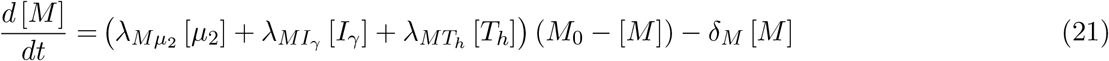

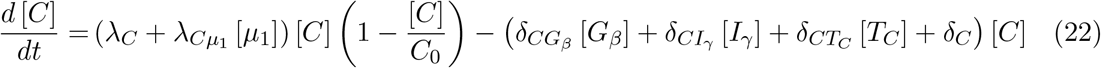

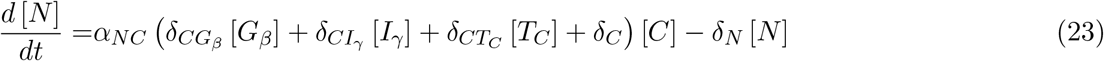

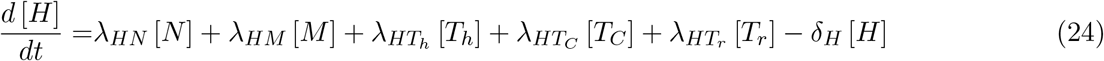

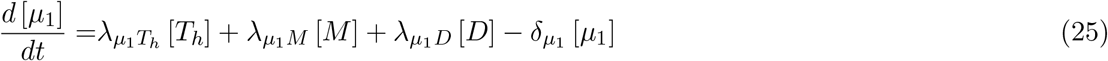

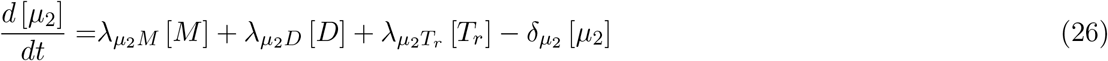

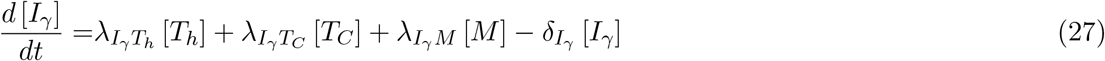

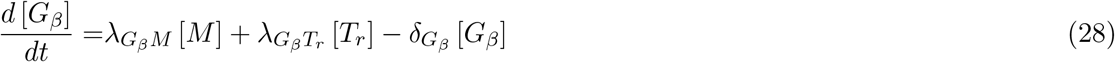

The system has 14 variables and 59 different parameters. The *λ* parameters correspond to proliferation, activation and production rates, *δ* parameters correspond to degradation and cell death rates, and four parameters: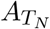 and 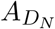 respectively are the production rates of naive T-cells and dendritic cells, *M*_0_ and *C*_0_ are the total density of macrophages (naive and activated together) and cancer cells maximum capacity, respectively.

### Appendix A.1 Positivity

To prove that the system with positive coefficients and positive initial conditions has positive solution let us consider the set of integrating factors, one for each variable:

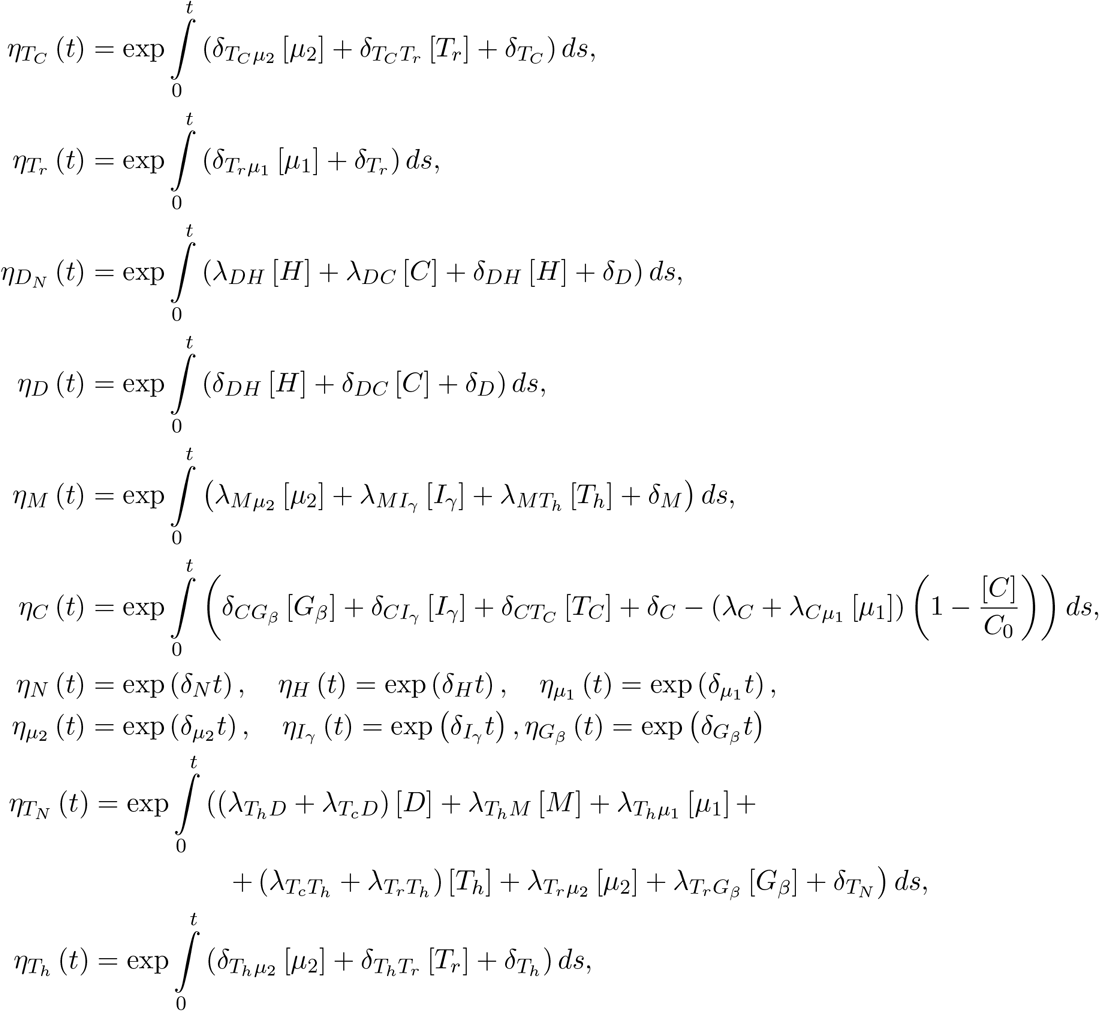

These integrating factors will not allow us to derive explicit solution as some of them are defined through the unknown variables. But it is important to note that the factors are strictly positive and allow us to rewrite the system as

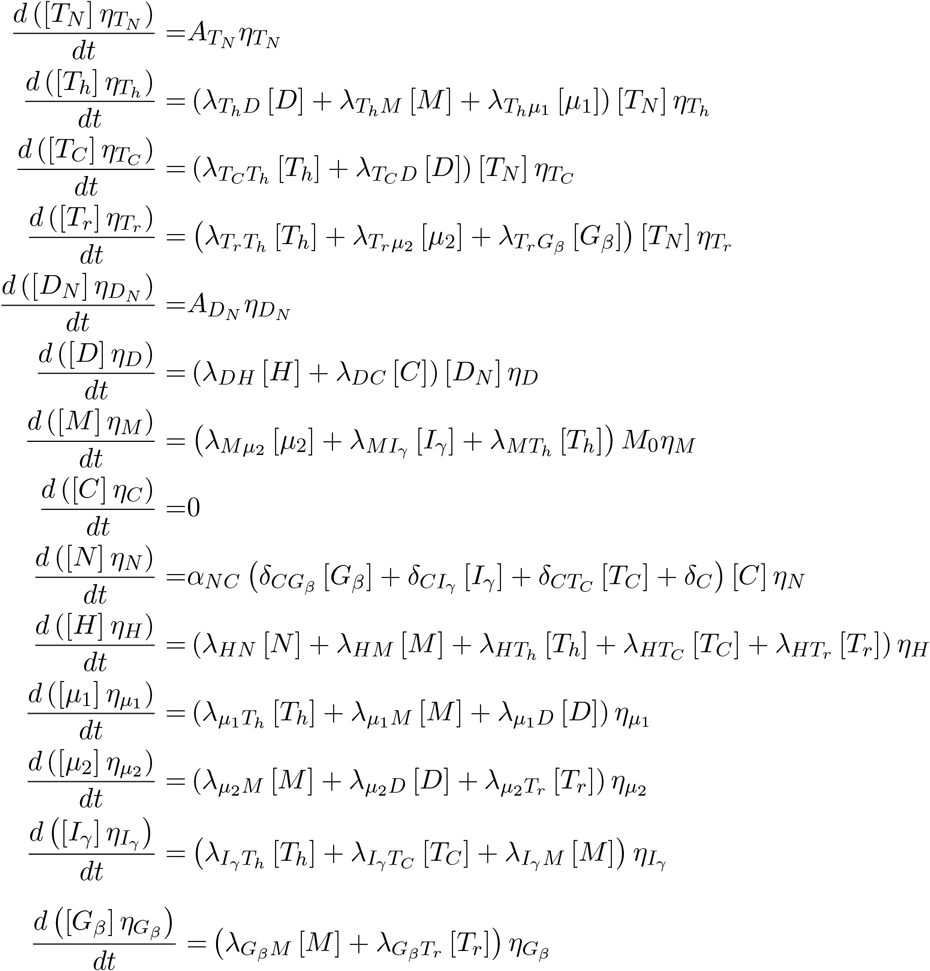

We see that the right-hand side of each equation in this system is non-negative, which means that the variable-factor product is non-decreasing, and thus if positive initially remains positive at all times.

### Appendix A.2 Boundedness

Let us show that all positive solutions are bounded for positive time *t*. We split the equations into groups by cell types. It is important to note, that we are not trying to derive the sharp bounds, just show that the bounds exist.

### T-cells

Adding equations (15)-(18) we get

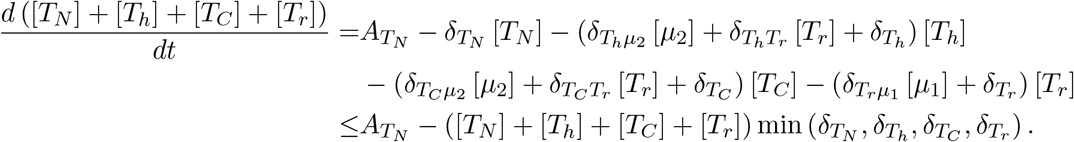

Integrating this inequality we obtain

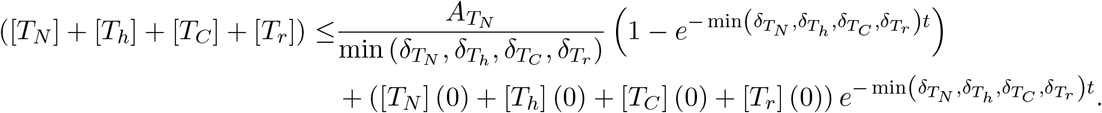

The right-hand side function is bounded, and since we have proven that each cell density is positive, all T-cells have to remain bounded.

### Dendritic cells

Let us add equations (19) and (20) to obtain

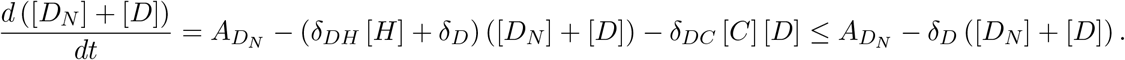

Integrating we get

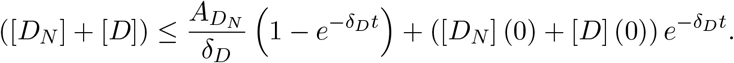

Since right-hand side is bounded and each variable is positive, this proves that each variable is bounded.

### Macrophages

Let us rewrite equation (21) as

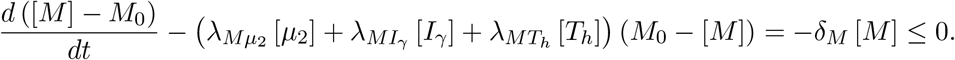

Integrating (with implicit dependence on variables [*µ*_2_], [*I*_*γ*_], and [*T*_*h*_]) results in

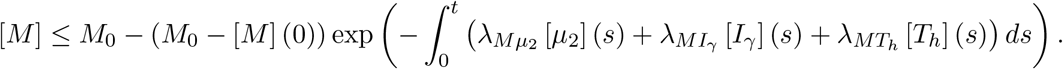

The right-hand side function is bounded for positive [*µ*_2_], [*I*_*γ*_], and [*T*_*h*_], and thus proves the bound on [*M*].

### Cancer cells

Rewriting the equation (22) as

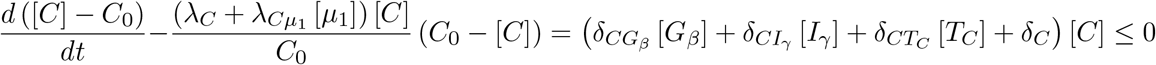

we integrate with implicit dependence on both [*C*] and [*µ*_1_] to obtain

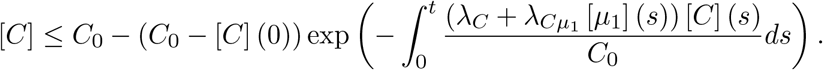

Since [*C*] and [*µ*_1_] are proven to remain positive, the right-hand side is bounded, hence [*C*] is bounded.

### Interferon-*γ* and TGF-*β*

Here we show the bound on [*I*_*γ*_] and [*G*_*β*_] as we need them to prove the bound on [*N*].

*Remark*. Alternatively we could show a bound on [*µ*_1_] and subsequent bound on [*N*] + [*C*]. Observe that in the right-hand sides of equations (27) and (28) the positive terms are already proven to be bounded:

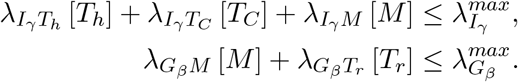

Then combining these with equations (27) and (28) we get the following differential inequalities:

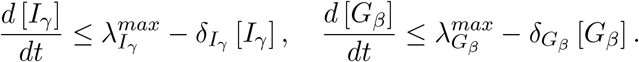

Integrating we get

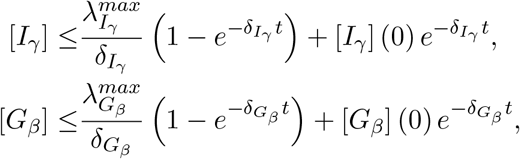

which proves the bound.

### Necrotic cells

Now we notice that for equation (23) all the components of the positive term are already proven to remain bounded

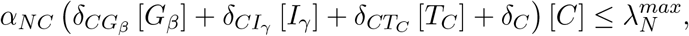

which results in the differential inequality

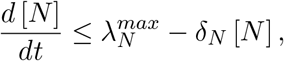

subsequently resulting after integration in the following bound:

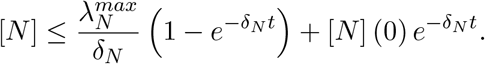

### Remaining cytokines

With the bound on necrotic cells we have proven boundedness for all the positive components of the right-hand sides of the equations (24)-(26):

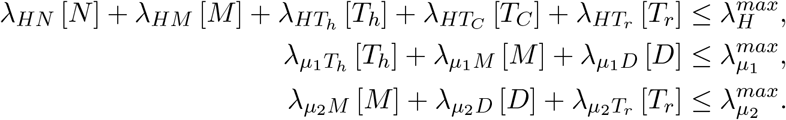

Thus the following differential inequalities are valid:

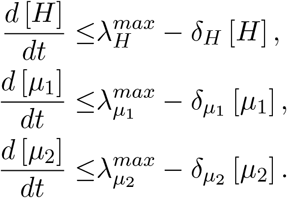

Integrating we obtain

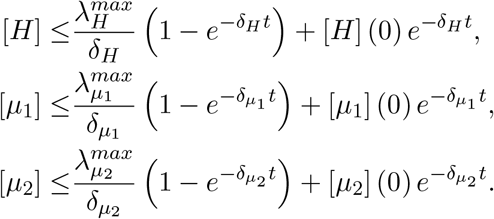

Thus [*H*], [*µ*_1_], and [*µ*_2_] are bounded for positive *t*.

## Appendix B Derivation of the sample parameter set

### Appendix B.1 Steady state and additional assumptions

We derive the sample parameter set under the assumption of specific values of steady state for each variable:

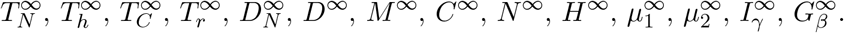

Then equations (15)-(28) provide us with 14 restrictions on parameters. There is a total of 59 parameters, so additional restrictions are required. We assume given cancer cell and macrophage capacities *C*_0_ and *M*_0_, as well as necrosis coefficient *α*_*NC*_ = 3*/*4. Additionally, from the available research [107, 125–134] we adopt the natural decay/death/degradation rates. For some of the specimen, considering a specimen *X* we will estimate death rate *δ*_*X*_ using published measurements of half-life 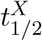 using the following formula 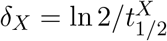 Other death rate estimates are provided directly in the referenced research. Here are the death rates used in our computations:

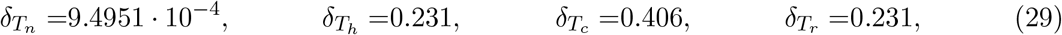

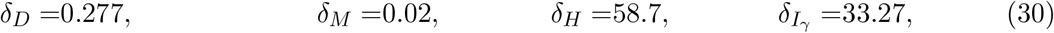

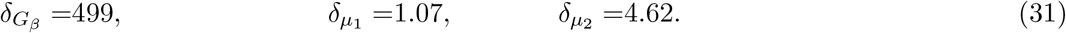

And as a last step, we impose heuristic assumptions on activation, inhibition and production rates. Let us look at these in detail.

First we consider the results in [135] suggesting that range of colon cancer doubling time is between 92.4 and 1032.2 days. Let us consider the doubling rate to be the difference between proliferation rate and death rate. Faster doubling rate includes both innate cancer proliferation and proliferation caused by *µ*_1_ family of cytokines, while death rate being only innate. This results in

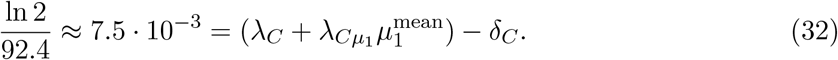

On the other hand for the slower doubling rate we only consider innate cancer proliferation, while death rate includes effects of all anti-cancer agents, i.e.

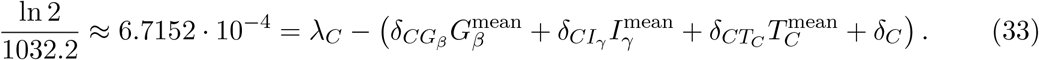

Here we consider 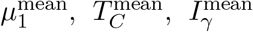, and 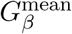 to be average values of the corresponding variable across all patients.

Further assumptions are based on maximal observable quantities for all the variable across all patients:

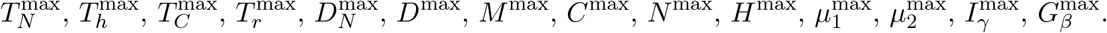

See Appendix C for more details on patient data and specific values.

We assume that most of T-helper cells are activated by antigen-presenting dendritic cells, so we take

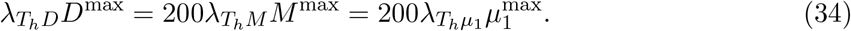

We also assume that inhibition of T-helper cells by *µ*_2_ family of cytokines and by Treg cells are each 20 times more effective than the natural degradation:

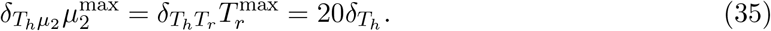

For cytotoxic T-cells we assume that activation by T-helper cells is twice as effective as activation by Dendritic cells

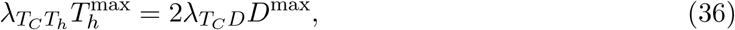

and same as for T-helper cells inhibition of cytotoxic T-cells by *µ*_2_ family of cytokines and by Treg cells are each 20 times more effective than the natural degradation:

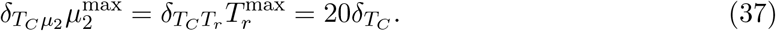

Next assumption is that activation of Treg cells by T-helper cells is four times larger than activation by *µ*_2_ family of cytokines and by TGF-*β*:

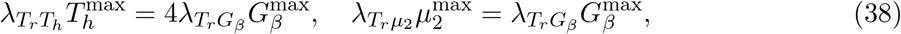

while inhibition of Treg cells by *µ*_1_ family of cytokines is 20 times larger than their natural degradation rate:

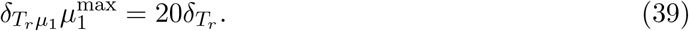

We impose that activation of dendritic cells by HMGB1 is twice more effective than activation by cancer cells, inhibition of dendritic cells by HMGB1 is twice less effective than inhibiiton by cancer cells, and cumulative inhibition of dendritic cells by HMGB1 and cancer cells is equivalent to the natural degradation rate of dendritic cells:

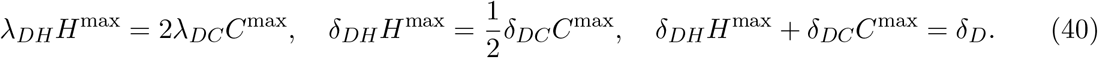

For macrophages we assume that most macrophages are activated by T-helper cells, thus

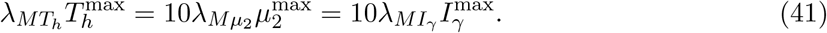

Next we look at the cancer death rates. We assume TGF-*β* and IFN-*γ* equally effective in killing cancer cells, but cytotoxic T-cells to be twice more effective, so

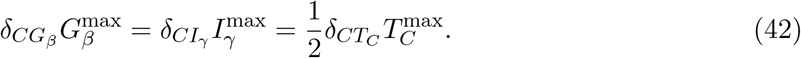

We also assume that at it’s extreme value TGF-*β* kills cancer cells 10 times faster than innate death rate of cancer cells:

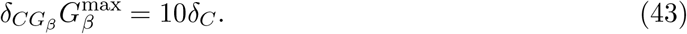

Next let us list assumptions on production rates of cytokines per cell:

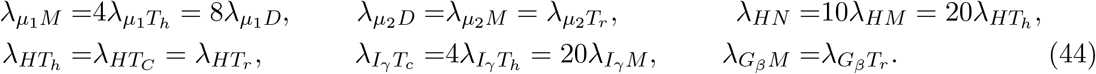

Altogether these assumptions are sufficient to derive a sample parameter set.

### Appendix B.2 Non-dimensionalization

For more stable numerical simulations and to avoid scale dependence in the sensitivity analysis, we introduce non-dimensional variables. For each variable [*X*] with steady state *X*^*∞*^ we introduce non-dimensional 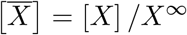 Then for all the non-dimensional variables steady state will be equal to 1. Because of the dramatic difference between timescales in different equations (related to natural decay rate) we make a choice to not scale time. Then we can rewrite the system as

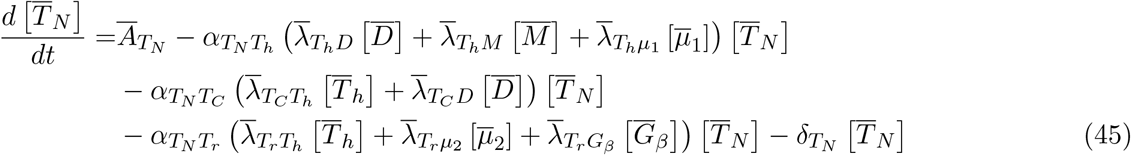

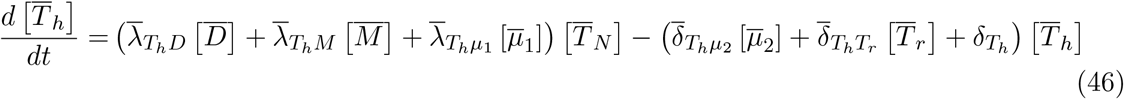

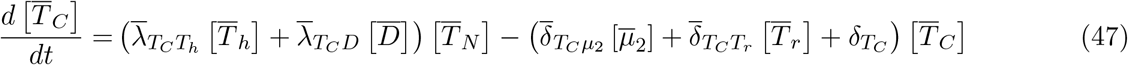

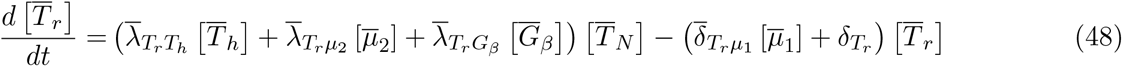

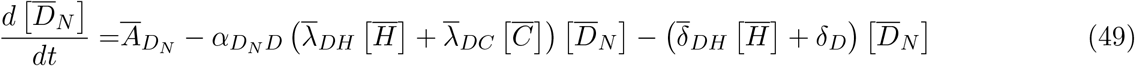

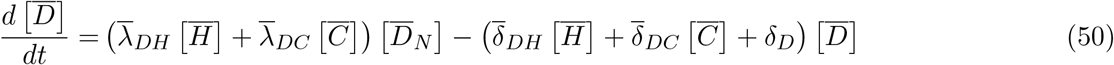

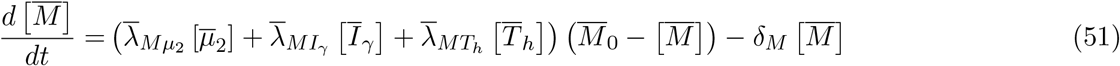

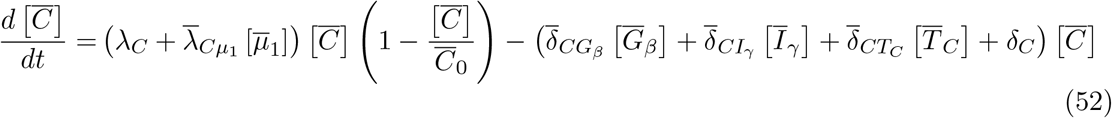

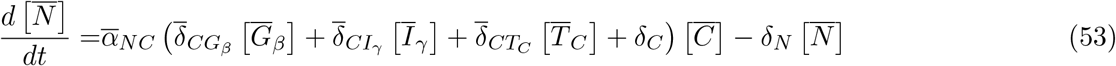

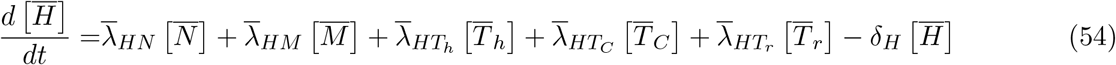

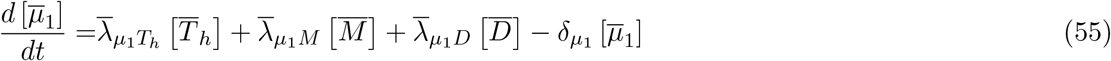

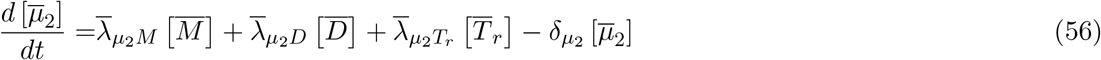

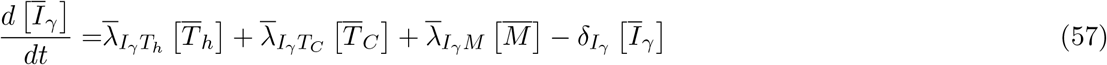

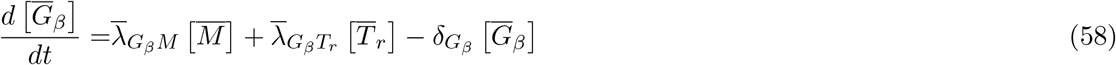

where the nondimensional parameters can be expressed as follows:

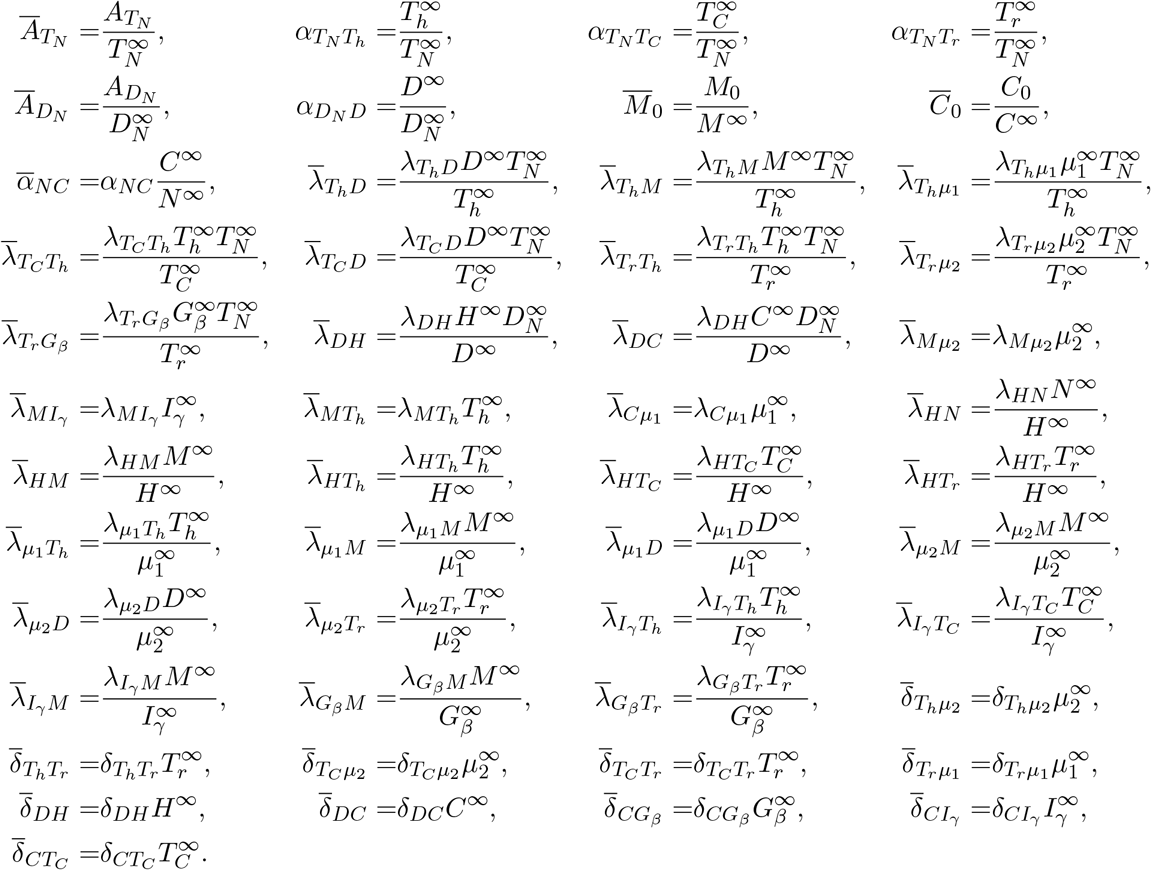

Cancer proliferation rate *λ*_*C*_ and all the innate degradation/death rates remain unscaled. Then the equations for doubling rate (32)-(33) become

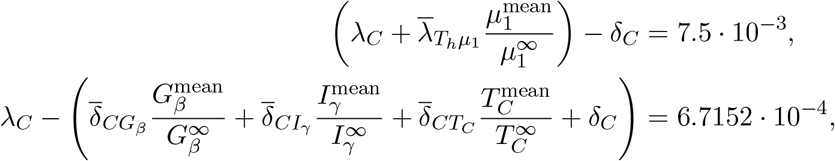

and the system of restrictions (34)-(44) in dimensionless form can be rewritten as

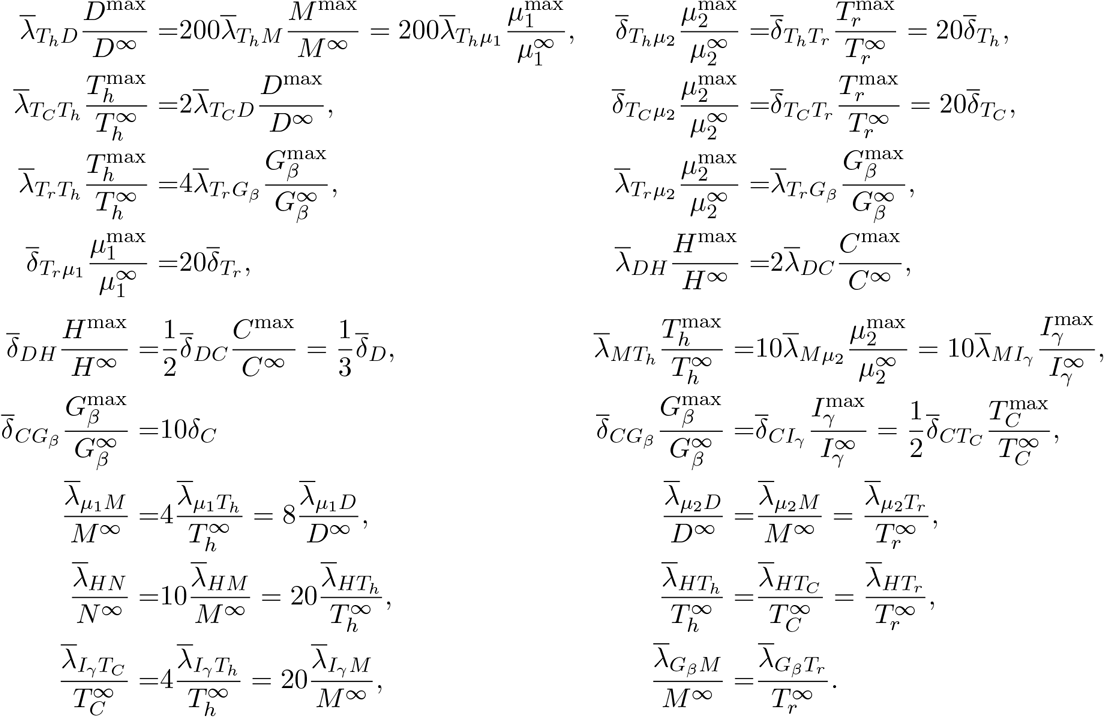

These 29 restriction, together with 14 equations from requiring steady state of (45)-(58), and 11 given decay rates (29)-(31), scaling coefficients

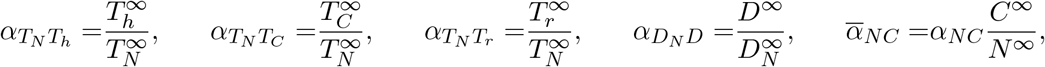

and given *α*_*NC*_, 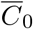 and 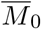 are enough to derive all 63 non-dimensional coefficients of (45)-(58) from

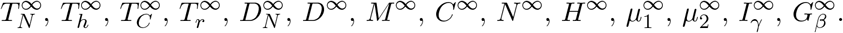

#### Appendix B.3 Parameter values

Here we detail all the parameter values derived and used in this paper. We divide them into three groups: innate degradation rates derived or adopted from prior research (see Table A1), scaling-independent parameters (those not affected by non-dimensionalization, see Table A2), and scaling-dependent parameters (see Table A3).

**Table A1:**
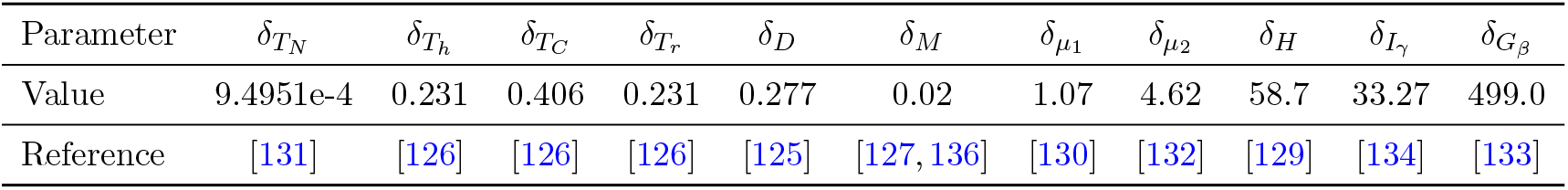
Prescribed parameters and their references. Innate degradation and death rates (in day^*−*1^) derived or adopted from given references.

| Parameter | $\delta_{T_N}$ | $\delta_{T_h}$ | $\delta_{T_C}$ | $\delta_{T_r}$ | $\delta_D$ | $\delta_M$ | $\delta_{\mu_1}$ | $\delta_{\mu_2}$ | $\delta_H$ | $\delta_{I_{\gamma}}$ | $\delta_{G_{\beta}}$ |
| --- | --- | --- | --- | --- | --- | --- | --- | --- | --- | --- | --- |
| Value | 9.4951e-4 | 0.231 | 0.406 | 0.231 | 0.277 | 0.02 | 1.07 | 4.62 | 58.7 | 33.27 | 499.0 |
| Reference | [131] | [126] | [126] | [126] | [125] | [127, 136] | [130] | [132] | [129] | [134] | [133] |

**Table A2:** Scaling-independent parameters. Values of scaling-independent parameters (in day^*−*1^) for each cluster derived from the steady state assumptions and patient data.

| parameter | Cluster 1 | Cluster 2 | Cluster 3 | Cluster 4 | Cluster 5 |
| --- | --- | --- | --- | --- | --- |
| $\lambda_C$ | 5.3323e-3 | 3.7596e-3 | 7.8327e-3 | 5.3535e-3 | 5.5150e-3 |
| $\delta_C$ | 7.8626e-4 | 5.2094e-4 | 1.2081e-3 | 7.8983e-4 | 8.1708e-4 |
| $\delta_N$ | 6.7332e-3 | 7.0593e-3 | 6.8602e-3 | 5.9142e-3 | 6.0514e-3 |

The scaling-independent parameters parameters include innate cancer proliferation rate *λ*_*C*_, innate cancer death rate (including apoptosis and necrosis) *δ*_*C*_, and necrotic cell degradation rate *δ*_*N*_. Because these parameters were not affected by non-dimensionalization procedure, as they are determined they remain independent of the scaling constant *α*_*dim*_, and depend solely on the derivation assumptions and patient data (thus different between clusters).

The scaling dependent parameters in their dimensional form in addition to derivation assumptions and patient data would also depend on the scaling constant *α*_*dim*_. Thus we prefer to list their more objective non-dimensional values. Because non-dimensionalization was done without time scaling, the dimension of most of these parameters is day^*−*1^. Exceptions are 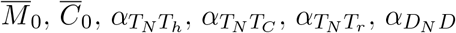, these are fully non-dimensional.

**Table A3:** Scaling-dependent parameters. Non-dimensional values of scaling-dependent parameters (in day^*−*1^, because the time was not scaled) for each cluster derived from the steady state assumptions and patient data.

| Parameter | Cluster 1 | Cluster 2 | Cluster 3 | Cluster 4 | Cluster 5 |
| --- | --- | --- | --- | --- | --- |
| $\bar{\lambda}_{T_h D}$ | 2.1399 | 2.7619 | 2.3163 | 1.7613 | 2.0179 |
| $\bar{\lambda}_{T_h M}$ | 3.9171e-2 | 4.3371e-2 | 7.8933e-2 | 3.4235e-2 | 1.1462e-1 |
| $\bar{\lambda}_{T_h \mu_1}$ | 1.1871e-2 | 1.9332e-2 | 5.0516e-2 | 6.6146e-3 | 2.6083e-2 |
| $\bar{\lambda}_{T_C T_h}$ | 3.2503 | 4.4295 | 3.6087 | 2.3377 | 3.5400 |
| $\bar{\lambda}_{T_C D}$ | 6.0053e-1 | 5.3505e-1 | 6.8998e-1 | 8.2970e-1 | 2.5392e-1 |
| $\bar{\lambda}_{T_r T_h}$ | 8.2898e-1 | 8.9260e-1 | 7.1048e-1 | 6.6821e-1 | 6.4634e-1 |
| $\bar{\lambda}_{T_r \mu_2}$ | 5.1120e-2 | 4.3090e-2 | 1.4558e-1 | 1.9891e-2 | 2.6898e-2 |
| $\bar{\lambda}_{T_r G_\beta}$ | 6.4120e-2 | 6.1990e-2 | 2.4271e-1 | 4.1296e-2 | 8.4474e-2 |
| $\bar{\lambda}_{DH}$ | 2.6134e-1 | 2.5410e-1 | 2.6749e-1 | 2.0531e-1 | 2.5253e-1 |
| $\bar{\lambda}_{DC}$ | 1.0129e-1 | 1.0998e-1 | 9.6275e-2 | 1.4619e-1 | 9.9406e-2 |
| $\bar{\lambda}_{M \mu_2}$ | 6.0389e-4 | 4.0918e-4 | 4.0357e-4 | 1.0688e-3 | 2.2488e-4 |
| $\bar{\lambda}_{M I_\gamma}$ | 5.4536e-4 | 4.4647e-4 | 3.5449e-5 | 5.5129e-4 | 2.4891e-4 |
| $\bar{\lambda}_{M T_h}$ | 2.4482e-2 | 2.1190e-2 | 4.9239e-3 | 8.9758e-2 | 1.3509e-2 |
| $\bar{\lambda}_{C \mu_1}$ | 3.6453e-3 | 5.6528e-3 | 1.3143e-3 | 2.5321e-3 | 2.5536e-3 |
| $\bar{\alpha}_{NC}$ | 1.5 | 1.5 | 1.5 | 1.5 | 1.5 |
| $\bar{\lambda}_{\mu_1 T_h}$ | 9.5154e-2 | 1.5848e-1 | 5.1746e-2 | 4.8884e-2 | 8.0520e-2 |
| $\bar{\lambda}_{\mu_1 M}$ | 9.6867e-1 | 9.0479e-1 | 1.0148 | 1.0150 | 9.8745e-1 |
| $\bar{\lambda}_{\mu_1 D}$ | 6.1798e-3 | 6.7287e-3 | 3.4777e-3 | 6.0984e-3 | 2.0302e-3 |
| $\bar{\lambda}_{\mu_2 M}$ | 3.6856 | 3.1869 | 3.2127 | 3.7737 | 3.7140 |
| $\bar{\lambda}_{\mu_2 D}$ | 1.8810e-1 | 1.8960e-1 | 8.8081e-2 | 1.8139e-1 | 6.1088e-2 |
| $\bar{\lambda}_{\mu_2 T_r}$ | 7.4634e-1 | 1.2435 | 1.3192 | 6.6490e-1 | 8.4487e-1 |
| $\bar{\lambda}_{HN}$ | 5.6645e+1 | 5.6824e+1 | 5.7494e+1 | 5.6720e+1 | 5.6504e+1 |
| $\bar{\lambda}_{HM}$ | 1.4603 | 1.0096 | 8.6817e-1 | 1.5130 | 1.6271 |
| $\bar{\lambda}_{HT_h}$ | 2.8689e-1 | 3.5366e-1 | 8.8539e-2 | 1.4573e-1 | 2.6536e-1 |
| $\bar{\lambda}_{HT_C}$ | 1.5995e-1 | 3.1527e-1 | 7.1133e-2 | 1.8821e-1 | 1.1817e-1 |
| $\bar{\lambda}_{HT_r}$ | 1.4786e-1 | 1.9697e-1 | 1.7824e-1 | 1.3328e-1 | 1.8507e-1 |
| $\bar{\lambda}_{I_\gamma T_h}$ | 8.8980 | 6.8580 | 6.4053 | 4.6181 | 9.8011 |
| $\bar{\lambda}_{I_\gamma T_C}$ | 1.9843e+1 | 2.4454e+1 | 2.0584e+1 | 2.3857e+1 | 1.7459e+1 |
| $\bar{\lambda}_{I_\gamma M}$ | 4.5290 | 1.9577 | 6.2806 | 4.7945 | 6.0098 |
| $\bar{\lambda}_{G_\beta M}$ | 4.1497e+2 | 3.5894e+2 | 3.5375e+2 | 4.2425e+2 | 4.0652e+2 |
| $\bar{\lambda}_{G_\beta T_r}$ | 8.4033e+1 | 1.4006e+2 | 1.4525e+2 | 7.4749e+1 | 9.2476e+1 |
| $\bar{\delta}_{T_h \mu_2}$ | 4.2911e-1 | 4.3775e-1 | 4.2642e-1 | 1.1131e-1 | 2.3641e-1 |
| $\bar{\delta}_{T_h T_r}$ | 1.5309 | 2.1559 | 1.7884 | 1.4599 | 1.6912 |
| $\bar{\delta}_{T_C \mu_2}$ | 7.5419e-1 | 7.6938e-1 | 7.4947e-1 | 1.9563e-1 | 4.1550e-1 |
| $\bar{\delta}_{T_C T_r}$ | 2.6906 | 3.7891 | 3.1432 | 2.5658 | 2.9724 |
| $\bar{\delta}_{T_r \mu_1}$ | 7.1322e-1 | 7.6668e-1 | 8.6777e-1 | 4.9839e-1 | 5.2671e-1 |
| $\bar{\delta}_{DH}$ | 3.3577e-2 | 3.1883e-2 | 3.5563e-2 | 1.9360e-2 | 2.9105e-2 |
| $\bar{\delta}_{DC}$ | 5.2053e-2 | 5.5200e-2 | 5.1199e-2 | 5.5139e-2 | 4.5828e-2 |
| $\bar{\delta}_{CG_\beta}$ | 9.1600e-4 | 7.1011e-4 | 1.8590e-3 | 3.9506e-4 | 1.3131e-3 |
| $\bar{\delta}_{CI_\gamma}$ | 6.5951e-4 | 5.3859e-4 | 9.7944e-5 | 9.8156e-5 | 4.6278e-4 |
| $\bar{\delta}_{CT_C}$ | 2.1270e-3 | 2.9365e-3 | 1.4085e-3 | 2.6597e-3 | 1.4414e-3 |
| $\bar{A}_{T_N}$ | 1.5006 | 4.1268 | 1.2183 | 1.1785 | 1.6150 |
| $\bar{A}_{Dn}$ | 1.0264 | 2.1172 | 3.5941e+2 | 1.8353 | 1.1435 |
| $\bar{M}_0$ | 1.7803 | 1.9072 | 4.7293 | 1.2189 | 2.4303 |
| $\bar{C}_0$ | 2.0 | 2.0 | 2.0 | 2.0 | 2.0 |
| $\alpha_{T_N T_h}$ | 3.1084e-1 | 5.2856e-1 | 1.5008e-1 | 1.7950e-1 | 3.6879e-1 |
| $\alpha_{T_N T_C}$ | 1.7330e-1 | 4.7118e-1 | 1.2057e-1 | 2.3182e-1 | 1.6423e-1 |
| $\alpha_{T_N T_r}$ | 1.6020e-1 | 2.9438e-1 | 3.0212e-1 | 1.6417e-1 | 2.5720e-1 |
| $\alpha_{D_N D}$ | 1.9739 | 4.9667 | 9.8717e+2 | 4.3783 | 2.3796 |

## Appendix C Patient data and initial conditions

Here we describe the processing of the data to be used for parameter estimation and initial conditions. The clustered deconvolution data, described in section 2.3, and original TCGA data is used to calculate the immune variables as described in table A4.

**Table A4:**
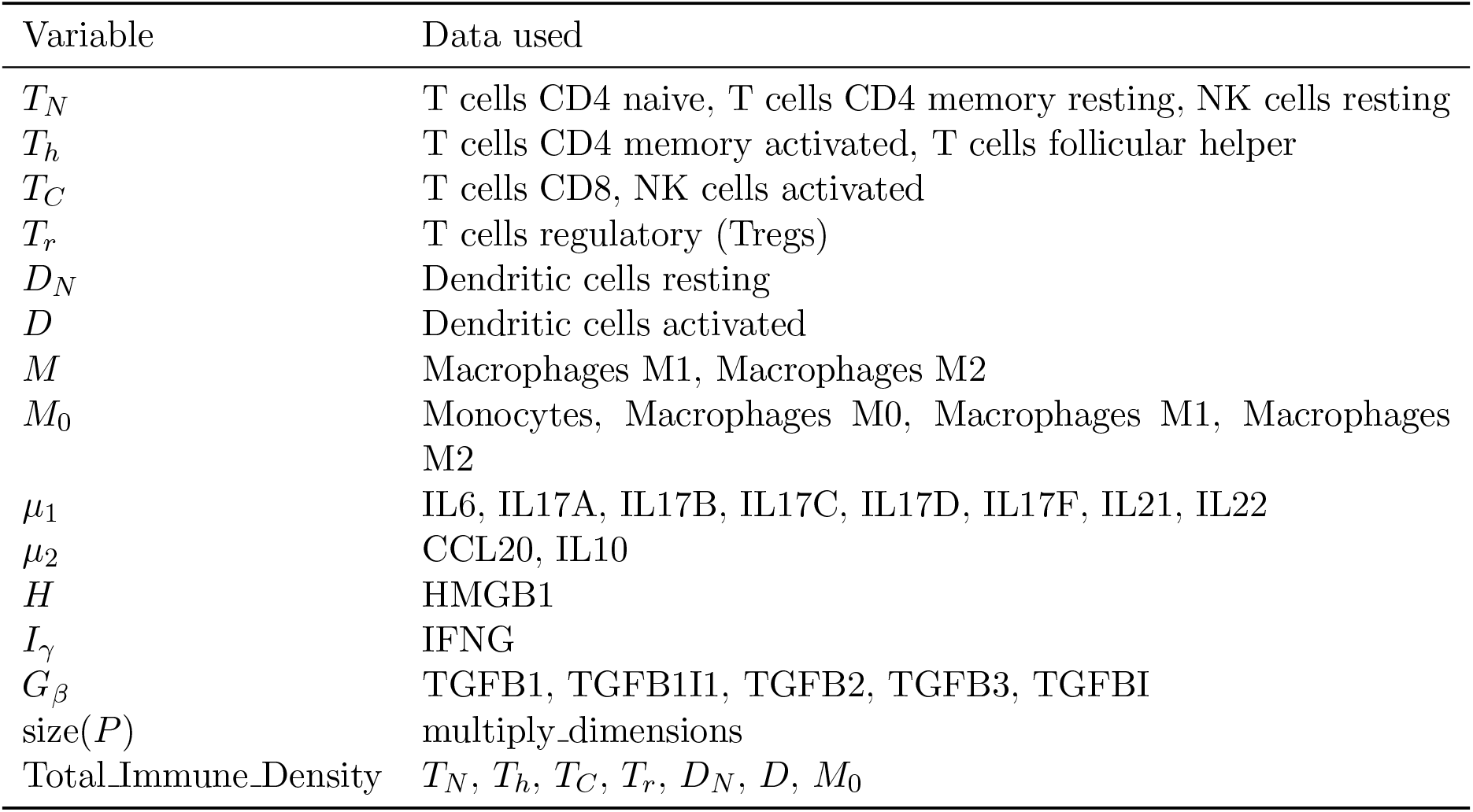
Patient data correspondence with variables. Correspondence between the system variables and the source data from TCGA and deconvolution results.

For variables related to immune cells we substitute zero values with 10% of the smallest positive cell density for numerical stability.

We estimate the relative amount of cancer cells and necrotic cells as follows: we start by assuming that on average across all patients the ratio of immune cells:cancer cells:necrotic cells will be approximately 0.4:0.4:0.2 with variability between clusters based on tumor size. For patient *P* we consider tumor size (size(*P*)) to be the product of the longest dimension and the shortest dimension. We assume total cell density at the steady state to be proportional to this product as

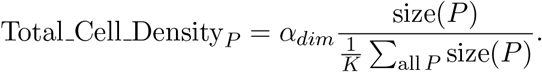

Tumor deconvolution data only provides us with ratios of immune cells relative to each other. Thus, to properly adjust the scaling, we take each immune cell value from deconvolution multiplied by 0.4*α*_*dim*_, and consider 0.4*α*_*dim*_ Σ(Immune cell ratios) as total immune density and

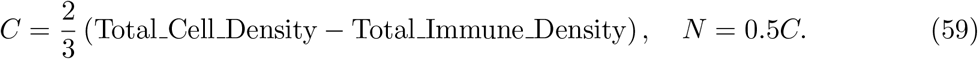

Next, for each cluster we divide patients into three groups according to their their tumor size: above average, below average and no data. Resulting patient numbers of each group are given in table A5. We use the means from the group “above average” as steady state assumptions given in table 2.

**Table A5:**
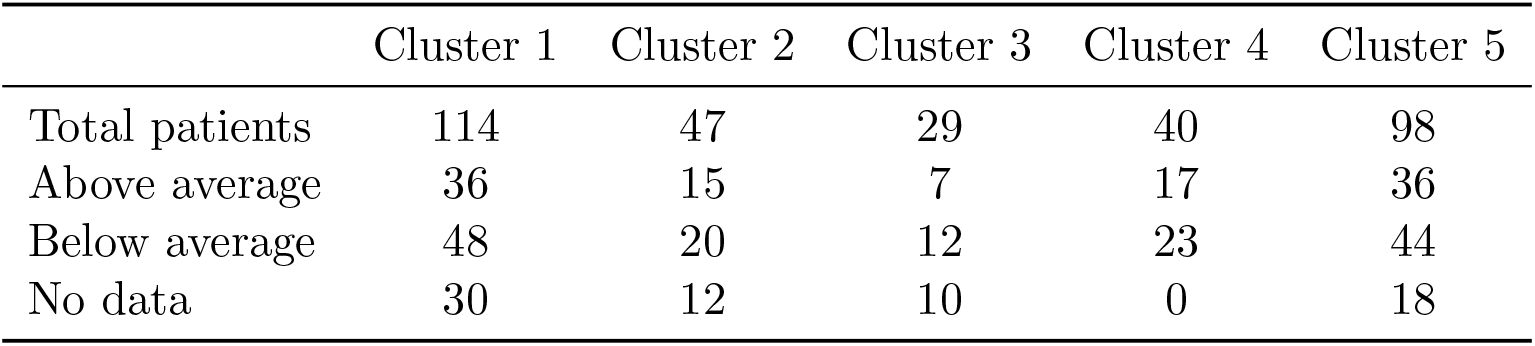
Distribution of patients according to their tumor size. Evaluated relative to the average tumor size within each cluster.

|  | Cluster 1 | Cluster 2 | Cluster 3 | Cluster 4 | Cluster 5 |
| --- | --- | --- | --- | --- | --- |
| Total patients | 114 | 47 | 29 | 40 | 98 |
| Above average | 36 | 15 | 7 | 17 | 36 |
| Below average | 48 | 20 | 12 | 23 | 44 |
| No data | 30 | 12 | 10 | 0 | 18 |

The data in the “below average” group as evaluated by (59) may contain negative values for cancer and necrotic cells. The data in “no data” group has no values for cancer and necrotic cells. Thus we substitute all non-positive and absent cancer values with 10% of the smallest positive cancer density value. We substitute all non-positive and absent necrotic cell values with zero. These changes violate the 0.4:0.4:0.2 ratio of immune cells:cancer cells:necrotic cells, and the updated ratio is 0.4475:0.3684:0.1841.

The steady state assumptions (see Appendix Appendix B) are partially based on maximum values of each variable in the ODE system (15)-(28) across all patients, as well as mean value of variables *T*_*C*_, *µ*_1_, *I*_*γ*_, and *G*_*β*_ across all patients. The corresponding values are given in table A6.

**Table A6:**
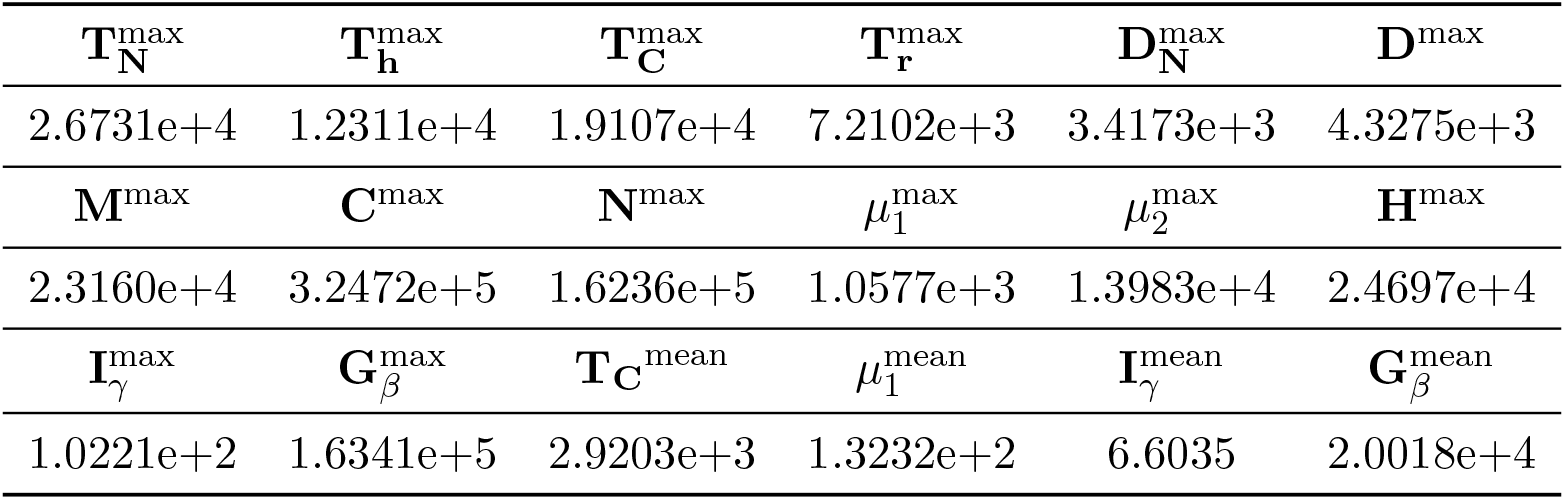
Maximum and mean variable values for parameter estimation. Maximum and mean cell densities in cells/cm^3^ and cytokine expression levels across all patients used in appendix Appendix B to derive parameter sets for time-dependent solutions.

In each cluster a patient with the smallest know tumor size is used as initial condition (given in table 3) for the dynamics computations presented in figures 4 and 5. However, any patient in the “below average” and “no data” group can be reasonable used as initial condition. The resulting dynamics is given by cluster on figures A1-A5.

**Figure A1:**
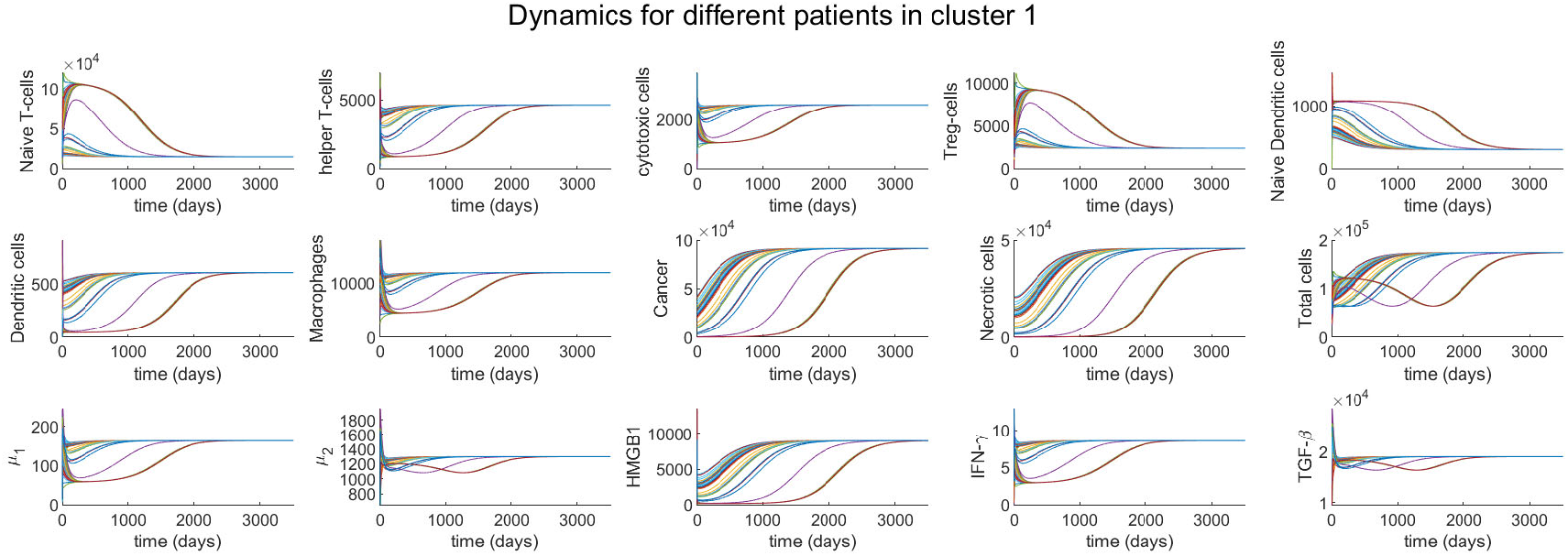
Different initial conditions for cluster 1. Based on patients in the “below average” and “no data” categories.

**Figure A2:**
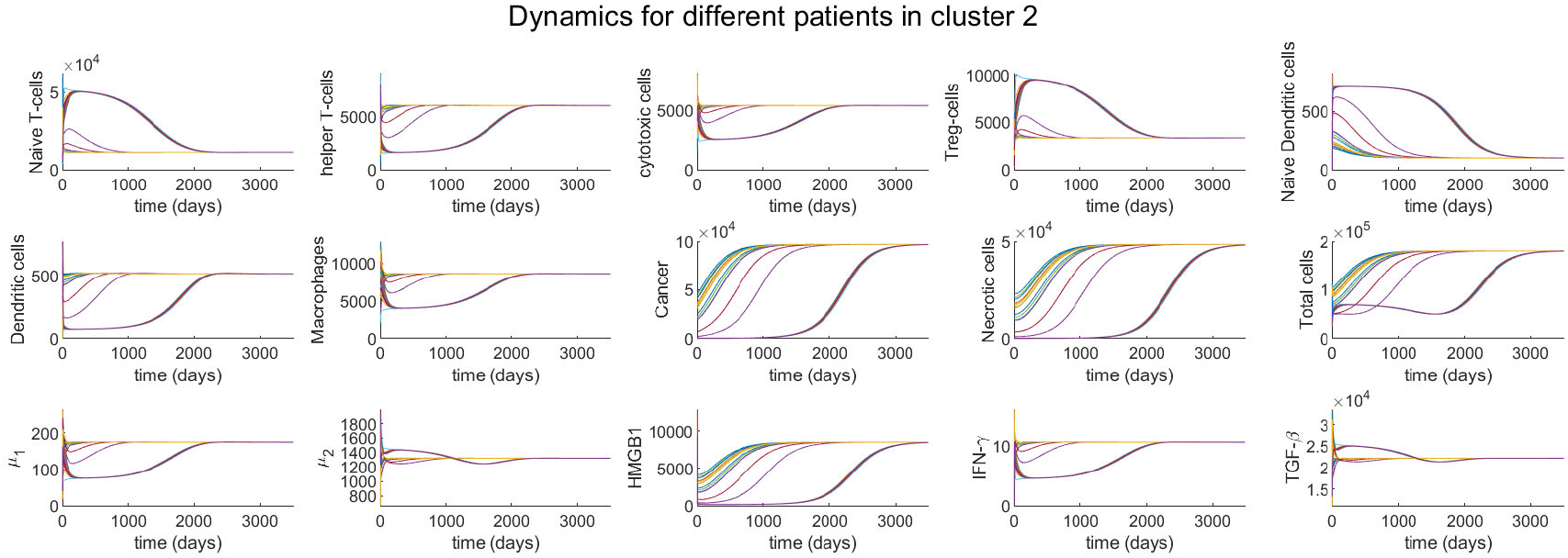
Different initial conditions for cluster 2. Based on patients in the “below average” and “no data” categories.

**Figure A3:**
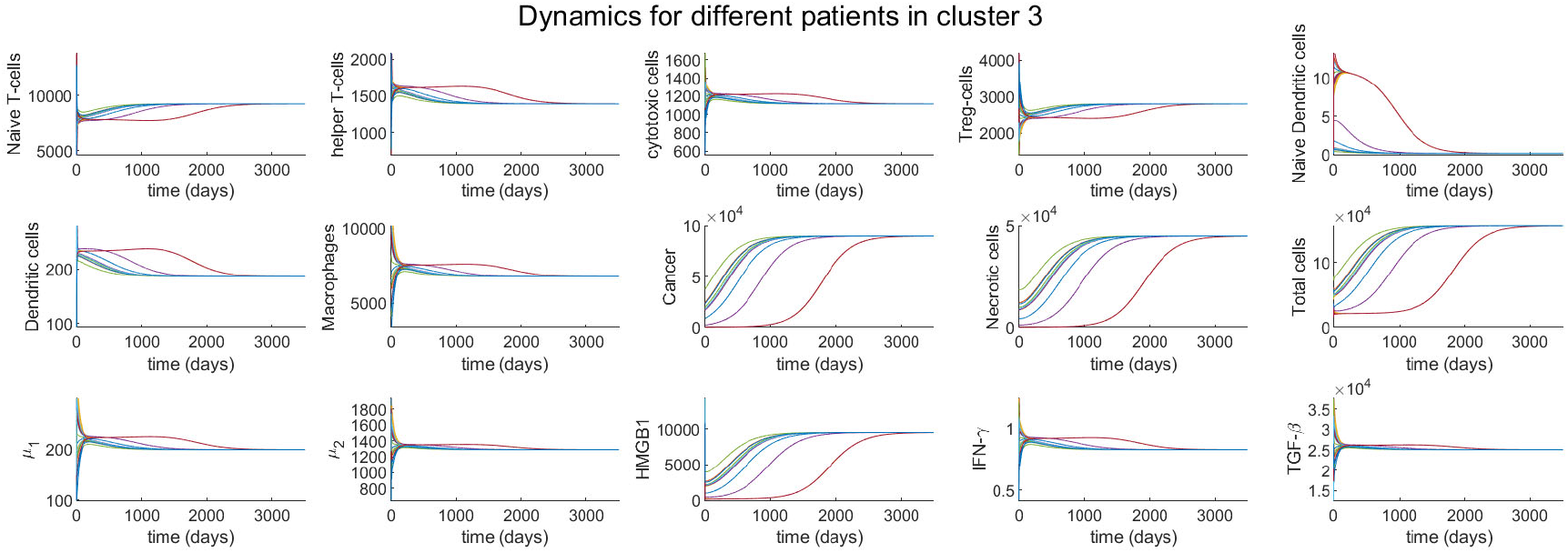
Different initial conditions for cluster 3. Based on patients in the “below average” and “no data” categories.

**Figure A4:**
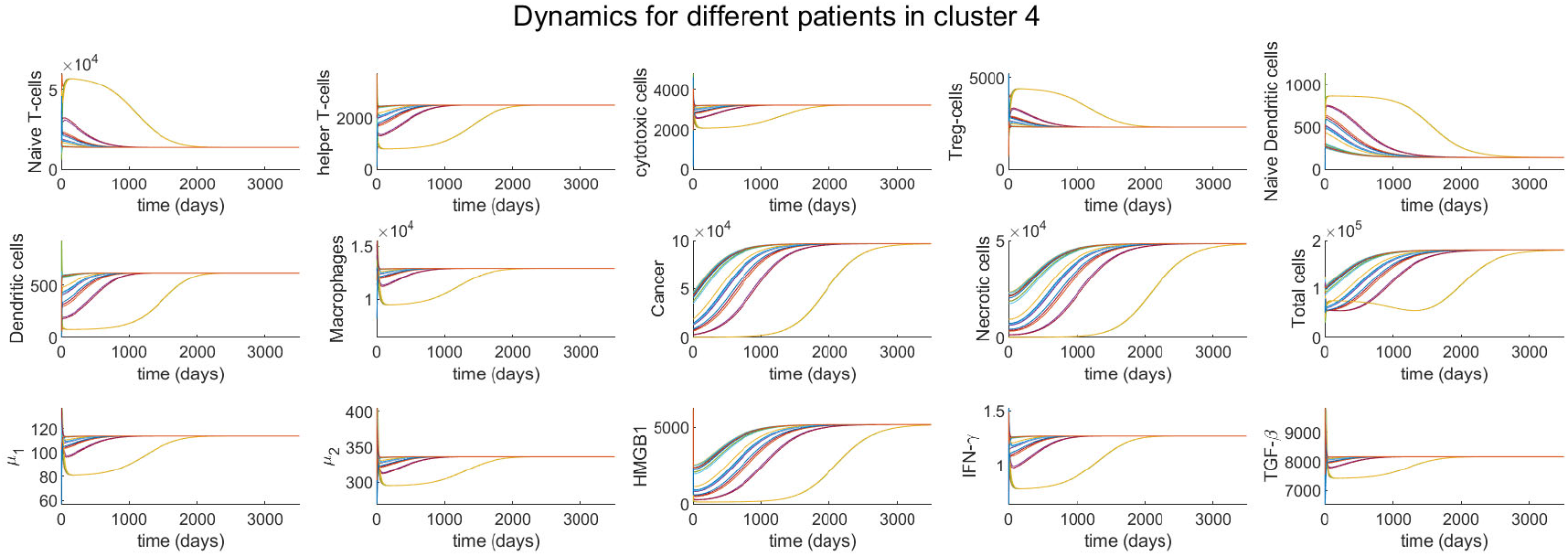
Different initial conditions for cluster 4. Based on patients in the “below average” and “no data” categories.

**Figure A5:**
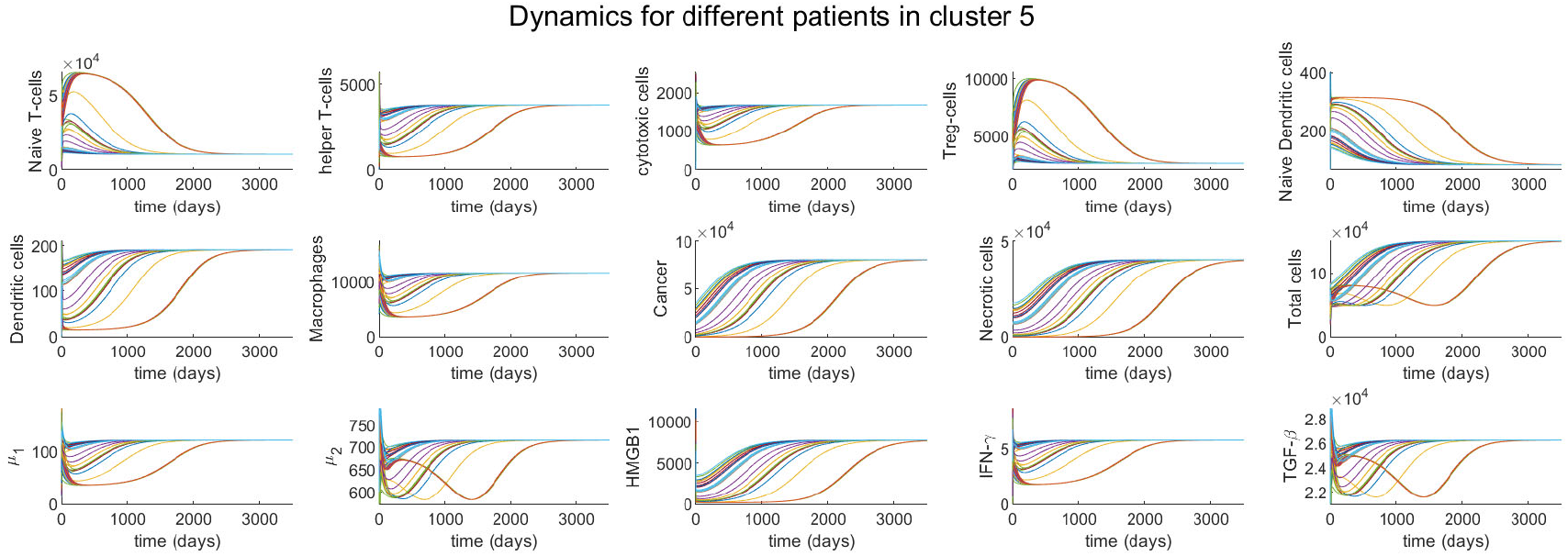
Different initial conditions for cluster 5. Based on patients in the “below average” and “no data” categories.

